# Selective Activation of GPCRs: Molecular Dynamics Shows Siponimod Binds but Fails to Activate S1PR2 Unlike S1PR1

**DOI:** 10.1101/2024.07.08.602194

**Authors:** Kumari Soniya, Kruthika Avadhani, Chanukya Nanduru, Antarip Halder

**Affiliations:** Computational Chemistry Department, Aganitha.ai, 512T, Road number 29, Jubilee Hills, Hyderabad, Telangana-500033, India

## Abstract

G Protein-Coupled Receptors (GPCRs) are central to drug discovery, accounting for nearly 40% of approved pharmaceuticals due to their regulatory role in diverse physiological processes. Given the high structural similarity among homologues, achieving receptor selectivity while minimizing off-target effects remains a major challenge in designing drugs targeting GPCRs. Sphingosine-1-phosphate receptors (S1PRs), comprising five subtypes, are therapeutically important GPCRs critical for immune and cardiovascular functions. Siponimod, an FDA-approved drug for multiple sclerosis, selectively modulates S1PR1 over S1PR2, unlike earlier S1PR modulators. However, the molecular basis for this selectivity is unclear, as cellular and biochemical assays provide limited insights. In this study, we used long-timescale molecular dynamics simulations to investigate how S1P and Siponimod binding affect S1PR1 and S1PR2 structural dynamics. Both ligands exhibited strong active site binding in both receptors. Crucially, while S1P and Siponimod induced similar activation-linked conformational changes in S1PR1, Siponimod failed to trigger these rearrangements in S1PR2. Specifically, Siponimod binding to S1PR2 led to altered side-chain dynamics of key TM7 residues (viz. Y^7.37^, F^7.38^, F^7.39^) and a drift of transmembrane helix 6 (TM6) towards orientations observed in inactive state. These unique structural features differentiate Siponimod’s behavior from S1P and explain its lack of inability to modulate S1PR2. Our findings elucidate molecular determinants of Siponimod’s selectivity towards S1PR1 and highlight these residues as potential differentiators for selective modulator design. This study demonstrates how structural and dynamic insights from atomistic simulations aid rational drug design for targets with high homology.

## Introduction

G protein-coupled receptors (GPCRs) constitute the most prevalent class of integral membrane proteins encoded in the human genome.^1^ Additionally, they comprise the largest family of membrane receptors that are the focus of approved pharmaceutical interventions. ^2^ However, designing a drug targeting GPCRs is challenging due to the high sequence and structural homology among members of the same sub-family, making it difficult to achieve selectivity. This similarity increases the risk of off-target effects, necessitating precise molecular design to differentiate between closely related receptors. ^3,4^

Among ∼800 GPCRs encoded in the human genome, five of the class A GPCRs are found to be highly specific towards sphingosine-1-phosphate (S1P) binding. ^5^ These are known as S1P receptors (S1PR1 through S1PR5). S1P, a signaling lysophospholipid, is produced by the phosphorylation of sphingosine via either sphingosine kinase 1 or sphingosine kinase 2.^6^ The S1P gradient regulates immune cell traffic between lymphoid organs and circulation through S1PR1.^7^ When S1PR1 binds to S1P, it undergoes internalization, reducing receptor presence on the cell surface and preventing lymphocytes from sensing the S1P gradient. As a result, lymphocytes are confined within lymphoid tissues.^8,9^ This mechanism makes S1PR1 a valuable target for treating autoimmune diseases such as Multiple Sclerosis (MS),^10–12^ tackling transplant rejections, and addressing conditions like Pulmonary Fibrosis.^11,13,14^ The receptor subtypes S1PR1, S1PR2 and S1PR3 are primarily expressed in the cardiovascular, central nervous, and immune systems, whereas the expression of S1PR4 and S1PR5 is restricted to the immune and nervous systems.^15,16^ S1PR2 is also linked with deafness in mice if the gene is mutated or knocked out.^17,18^

Certainly, while designing drugs for S1PR1, it is ideal to ensure that the drugs do not modulate other receptors, specifically S1PR2 and S1PR3 due to their involvement with cardiac system. For example, the first-in-class approved drug for MS, Fingolimod (FTY720), is known to be active against all S1P receptors except S1PR2.^3,19^ Activation of S1PR3 by Fingolimod is associated with heart conditions such as bradycardia. ^20,21^ Hence, this limits the clinical use of Fingolimod. In contrast, second-generation S1P receptor modulators such as Siponimod, Ozanimod, and Ponesimod are more selective for S1PR1 and S1PR5 over S1PR2, S1PR3, and S1PR4, resulting in fewer side effects. ^9,11,12^ Understanding the mechanisms of action of these next-generation drugs is crucial for developing strategies to enhance drug selectivity in GPCR therapeutics.

Functional assays, such as the GTP*γ*[^35^S]-binding assay, are commonly employed to assess the activity of ligands against GPCRs.^22,23^ When agonists bind to GPCRs, they induce conformational changes in the receptor, leading to the dissociation of the *α* subunit from the *β* and *γ* subunits of the heterotrimeric G protein complex.^24^ This assay measures the radioactivity of GTP*γ*[^35^S] that binds to the G*α* subunit by displacing GDP from the binding pocket. However, these assays cannot differentiate whether G-protein activation failed due to the ligand not binding to the active site or failing to induce the necessary structural changes despite binding. Siponimod is a second-generation approved treatment in adults for the relapsing forms of Multiple Sclerosis (MS) that includes the clinically isolated syndrome, relapsing-remitting MS, and active secondary progressive MS. ^25,26^ *In vitro* studies using GTP*γ*[^35^S]-binding assay reported that the half maximal effective concentrations (EC_50_) for Siponimod in human S1PR1 and S1PR5 are ≈0.46 nM and ≈ 0.3 nM, respectively. In contrast, the EC_50_ for S1PR2, S1PR3 and S1PR4 are > 10,000 nM, > 1111 nM and ≈383.7 nM, respectively.^22,27^ So, it is evident that Siponimod selectively modulates S1PR1 and S1PR5 over S1PR2, S1PR3, and S1PR4. Also, radioligand binding assays confirm Siponimod’s binding at the active site (i.e., the S1P binding site) of human S1PR1 and S1PR5 but the results for the same are not available for the other three receptors that Siponimod fails to modulate.^27^

Meticulously designed computational studies have the potential to complement experimental methods and help overcome their limitations, particularly in the context of drug design.^28^ For example, a recent study integrating various computational protocols like blind docking, conventional as well as enhanced molecular dynamics simulations etc. revealed a significantly lower activation barrier and a more stable intermediate state for Siponimod activation of S1PR1 compared to S1PR3.^29^ In this study, we used a range of bioinformatics and computational chemistry techniques to explore the potential binding of Siponimod to the active site of S1PR1 and S1PR2. Previous docking approaches suggested that Siponimod’s lower selectivity for S1PR2 might be due to steric clashes with a phenylalanine residue (F274) upon docking to its active site.^30^ However, this hypothesis was based on an engineered active site of S1PR2 derived from a cryo-EM structure of Siponimod-bound S1PR1. Recently, the cryo-EM structure of S1P-bound S1PR2 has been resolved,^17^ providing an opportunity to investigate Siponimod’s binding potential to S1PR2 using more accurate computational methods. Our analyses, which include structure modeling, molecular docking, microsecond-timescale atomistic Molecular Dynamics (MD) simulations, and enhanced MD simulations, demonstrate that Siponimod binds effectively to the active site of S1PR2. However, despite this strong binding affinity, Siponimod fails to trigger the conformational changes required for activation of S1PR2. Based on these computational insights, we hypothesize that Siponimod’s inability to activate S1PR2 stems from the absence of critical interactions with specific key residues—interactions that are present in its binding to S1PR1. Identification and consideration of these key residues are therefore crucial for guiding the design of selective modulators that can discriminate between receptor subtypes by modulating activation rather than binding alone.

## Materials and methods

Sequence alignment of S1PR1 and S1PR2 was performed using NCBI BLASTp^31,32^ server to determine the percentage sequence similarity between the two S1P receptors.

### Structure modelling of S1PR1 and S1PR2

The general architecture of the seven transmembrane helices along with extracellular and intracellular loops in S1PRs are illustrated in Figure 1a. Important transmission switches, ERY(H) and NPxxY motifs along with most conserved residues as per Ballesteros Weinstein (BW)^33^ numbering schemes are depicted in the snake plot.^34,35^ For *in silico* investigations, we selected the latest cryo-EM structures of human S1PR1 (PDB ID: 7TD3^30^) and S1PR2 (PDB ID: 7T6B^17^) that are complexed with the native ligand S1P and with the G proteins in the cytoplasmic regions. This will ensure that the S1P receptors are in their respective active state conformations. For the next steps we focused only on the S1PR chains, i.e. chain D in 7TD3 and chain E in 7T6B. Missing residues in both the chains, except the highly disordered N and C terminal residues, were modelled using Swiss modeler server. ^36^ The final modeled S1PR1 and S1PR2 chains have 304 and 279 residues, respectively. The first 21 residues in the N-terminal are missing in final modelled S1PR1 and the first 11 N-terminal residues are missing in the final modelled S1PR2. The residue numbering was adjusted accordingly. The structural similarity between the final modeled structures of S1PR1 and S1PR2 was assessed using Foldseek.^37^ Foldseek performs structural alignment by utilizing a structural alphabet that emphasizes tertiary interactions rather than the protein backbone. This method generates a TM-score^38^ to quantify structural similarity, ranging from 0 to 1. A higher TM-score indicates greater structural similarity between the two proteins.

**Figure 1:**
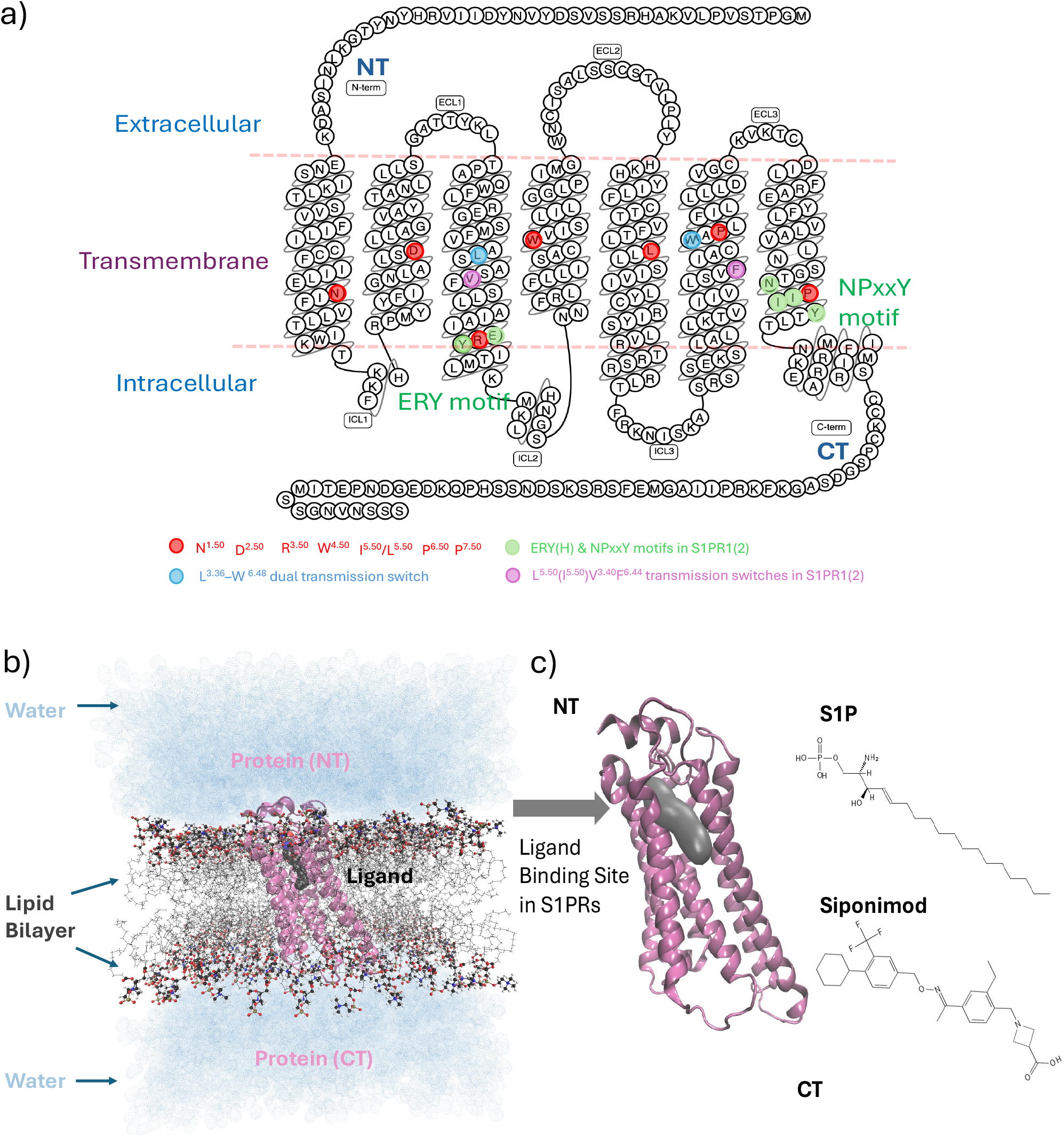
System details for S1P receptors. a) Snake plot (using S1PR1 as reference) depicting the most conserved residues of the transmembrane (TM) helices in S1PRs (red), important toggle switches (blue) and conserved ERY(H) and NPxxY motifs (green) in S1PRs; b) MD simulation system consisting of ligand bound S1P receptors inside lipid bilayer in aqueous medium; c) S1P and Siponimod binds towards the N-terminal (NT) end of the receptors occupying same binding pocket as shown in gray surface.

### Molecular docking

Molecular docking was performed using AutoDock VINA^39,40^ utility in UCSF Chimera^41^ with default settings. S1P and Siponimod were docked at the S1P binding site (illustrated in Figure 1c) of both S1PR1 and S1PR2. Different non-covalent interactions forming between the docked ligand and the binding site residues were analyzed using Protein-Ligand Interaction Profiler (PLIP).^42^ The docked poses with the highest docking scores were used for building the initial structures for performing MD simulations. To mimic the cellular environment in the MD simulations, ligand free and ligand bound receptors were embedded in lipid environment and surrounded by water as illustrated in Figure 1b. All-atom unbiased MD simulations were performed on six systems, viz., (i) Siponimod bound S1PR1 (Siponimod-S1PR1), (ii) Siponimod bound S1PR2 (Siponimod-S1PR2), S1P bound S1PR1 (S1P-S1PR1), (iv) S1P bound S1PR2 (S1P-S1PR2), (v) ligand free S1PR1 (apo S1PR1) and (vi) ligand free S1PR2 (apo S1PR2).

### Setting up MD simulation

The MD systems were prepared using a set of utilities present in AmberTools22.^43^ Force fields for S1P and Siponimod were generated with AM1-BCC charges^44,45^ in antechamber.^46,47^ Protein was embedded in the POPC (1-palmitoyl-2-oleoyl-sn-glycero-3-phosphocholine) layer with proper alignment using PACKMOL-Memgen ^48^ tool in AmberTools22. Lipid21 force field^49^ was used for the lipid bilayer. The systems were hydrated with a three-point SPC/E^50,51^ water model in a cubic box with 8 Å buffer. Chloride ions were added to neutralize the system. The system was then minimized using sander engine on single CPU. Minimization was carried out using steepest descent algorithm for the first 25,000 cycles, and conjugate gradient algorithm was used for last 25,000 cycles. AMBER minimized topology and coordinate files of the systems were converted to GRO-MACS compatible format using ParmEd python utility. Minimization, NVT equilibration followed by NPT equilibration and NPT production run were carried out in GPU enabled GROMACS 2022.3^52–54^ following well-established protocols as discussed in detail in the SI. For each of the six systems, ≈1*µ*s long MD trajectory was generated for further analysis. For the four ligand bound cases (i-iv), a second replica of ≈1*µ*s was run for statistical validation of the observed conformational differences. For all ligand-bound systems, we conducted an additional five ≈20 ns simulations. This was done to obtain more robust statistical estimates of binding free energies and residence times, as discussed further in the text. For both the S1P-bound and Siponimod-bound S1PR2 systems, we also performed additional ≈500 ns MD simulations. These longer runs were crucial for validating the conformational and structural change insights derived from our initial two ≈1*µ*s trajectories.

### Analysing MD trajectories

The MD trajectories were visualized using Visual Molecular Dynamics (VMD 1.9.4) tool^55^ and analyzed using GROMACS and AmberTools22. Fluctuations in receptors during the simulation trajectories were measured by Root Mean Square Displacement (RMSD) with respect to the initial structure and Root Mean Square Fluctuations (RMSF) averaged over the entire trajectory. All RMSD values were computed using GROMACS by aligning trajectories on the protein backbone atoms and calculating RMSD over the same backbone atom set. An array of collective variables (CVs) were used to track the structural changes in the transmission switches and mutual motions of transmembrane (TM) helices. Details of those customized CVs are given in the Table SI 1 of Supporting Information. To compare the relative dynamics of different parts of the receptors, Dynamic Cross Correlation Maps (DCCM) and Principal Component Analysis (PCA)^56^ were obtained using CPPTRAJ utility^57^ from AmberTools22.

To build a Markov State Model (MSM),^58^ Principal Component Analysis (PCA) was first applied to the atomistic MD trajectories to reduce dimensionality. All the heavy atoms were considered for the PCA. The two main components (PC1 and PC2) that collectively explained at least 80% of the total variance were selected. The MD trajectory was then projected onto these selected PCs, creating a lower-dimensional representation of the conformational landscape. Subsequently, these projected data were clustered using a k-means algorithm to define a set of discrete metastable states. Finally, a transition matrix was estimated from the transitions between these states over a chosen lag time, allowing for the construction and validation of the MSM to characterize the system’s convergence. PyEMMA^59^ package was used to cluster and build MSM.

Relative binding free energies of the ligand-bound receptors were obtained using MMG-BSA calculations^60^ implemented through the `gmx_MMPBSA` package.^61^ A single-trajectory approach based on a modified Gibbs-Boltzmann protocol was employed. ^62^ A salt concentration of 0.15 M was used for all calculations. To better understand the chemistry of ligand–protein interactions, pair-wise energy decomposition analysis was carried out using the `idecomp=3` setting, which includes the 1–4 interactions in the internal energy terms. Residues within 4Å of the ligand were selected for detailed analysis of their individual contributions to the binding free energy. The step-wise procedure for consolidating data from two independent replicas is provided in the Supporting Information.

### *τ*RAMD

*τ*RAMD^63^ simulations were performed to estimate ligand residence times using the latest gromacs-RAMD implementation (gromacs-ramd-gromacs-2024.1-ramd-2.1) available at https://github.com/HITS-MCM/gromacs-ramd. For each of the five replicas used in the MMGBSA binding affinity calculations, 20 RAMD trajectories were generated with different random seeds. A random force of 14.35 kcal/mol was applied to the ligand, and dissociation was tracked until a maximum displacement of 30 Å from the binding pocket. The RAMD force was evaluated every 50 integration steps. Final *τ*RAMD values were computed as the mean residence time across all 20 trajectories from each of the five replicas.

## Results and discussion

This study explores the selective modulation of S1PR1 over S1PR2 by Siponimod using a comprehensive array of *in silico* methods, including protein sequence and structure alignment, molecular docking, and all-atom conventional MD simulations. The BW numbering scheme is utilized in this article to identify the residues in the transmembrane helices.

### Active sites of both S1PR1 and S1PR2 have high structural similarity

The sequence and structural similarities between the two S1P receptors, S1PR1 and S1PR2, were analyzed in detail. A sequence alignment using the NCBI BLASTp server reveals that these two sequences share only about 50% similarity (Figure SI 1). Despite this modest sequence similarity, the proteins exhibit high structural similarity, as evidenced by a TM-score of 0.88 obtained from Foldseek (Figure SI 2). This is expected since both receptors are members of the S1P receptor sub-family. The transmembrane helices and the overall fold, particularly the configuration of the S1P binding pocket (the active site), are nearly identical in both receptors. Minor structural differences are primarily due to variations in the orientations of the extracellular (ECL) and intracellular (ICL) loops (Figure SI 2). Given the structural similarities of the active sites in S1PR1 and S1PR2 and considering that Siponimod binds to the active site of S1PR1, it is reasonable to hypothesize that Siponimod will also bind to the active site of S1PR2.

While most residues involved in ligand binding and activation mechanisms are conserved, the sequence alignment indicates a few notable differences (Figure SI 1). For example, leucine (L^7.39^) in S1PR1 is replaced by phenylalanine (F274/F^7.39^) in S1PR2, highlighted in pink in Figure SI 1. Other similar differences are discussed in the SI. Although both leucine and phenylalanine have hydrophobic side chains, phenylalanine’s bulkier phenyl group was previously hypothesized^30^ to cause a steric clash, preventing Siponimod from binding to the active site of S1PR2 based on studies using an engineered binding pocket. However, with the availability of the S1P-bound S1PR2 structure,^17^ we can now re-evaluate this hypothesis and determine whether Siponimod can indeed bind to the active site of S1PR2.

### Molecular docking suggests Siponimod can bind strongly to the active sites of both S1PR1 and S1PR2

We utilized molecular docking to compare the binding affinities of both S1P and Siponimod to the active sites of S1PR1 and S1PR2. At the outset of the discussion, it is important to note that experimentally determined structures of S1P-bound S1PR1 and S1PR2 were used in this study. The apo and Siponimod-bound systems for S1PR1 and S1PR2 were prepared from these S1P-bound protein structures, suggesting that the initial structures of S1PR1 and S1PR2 were already in an active conformation. Siponimod was then docked to the active conformations of both receptors. Both S1P and Siponimod occupy the same space inside the active site with polar head facing the extracellular region and the non-polar tails buried in the hydrophobic pocket near the transmission switches. The binding affinity of Siponimod for S1PR2 (−13 kcal/mol) was ≈1.5 kcal/mol higher than that for S1PR1 (−11.7 kcal/mol). Additionally, contrary to the previous hypothesis,^30^ the docked pose of Siponimod showed no steric clashes with F^7.39^ (F274) at the S1PR2 binding site (Figure 2c). These findings indicate that Siponimod not only binds to both receptors but also exhibits a slightly stronger binding affinity for S1PR2.

**Figure 2:**
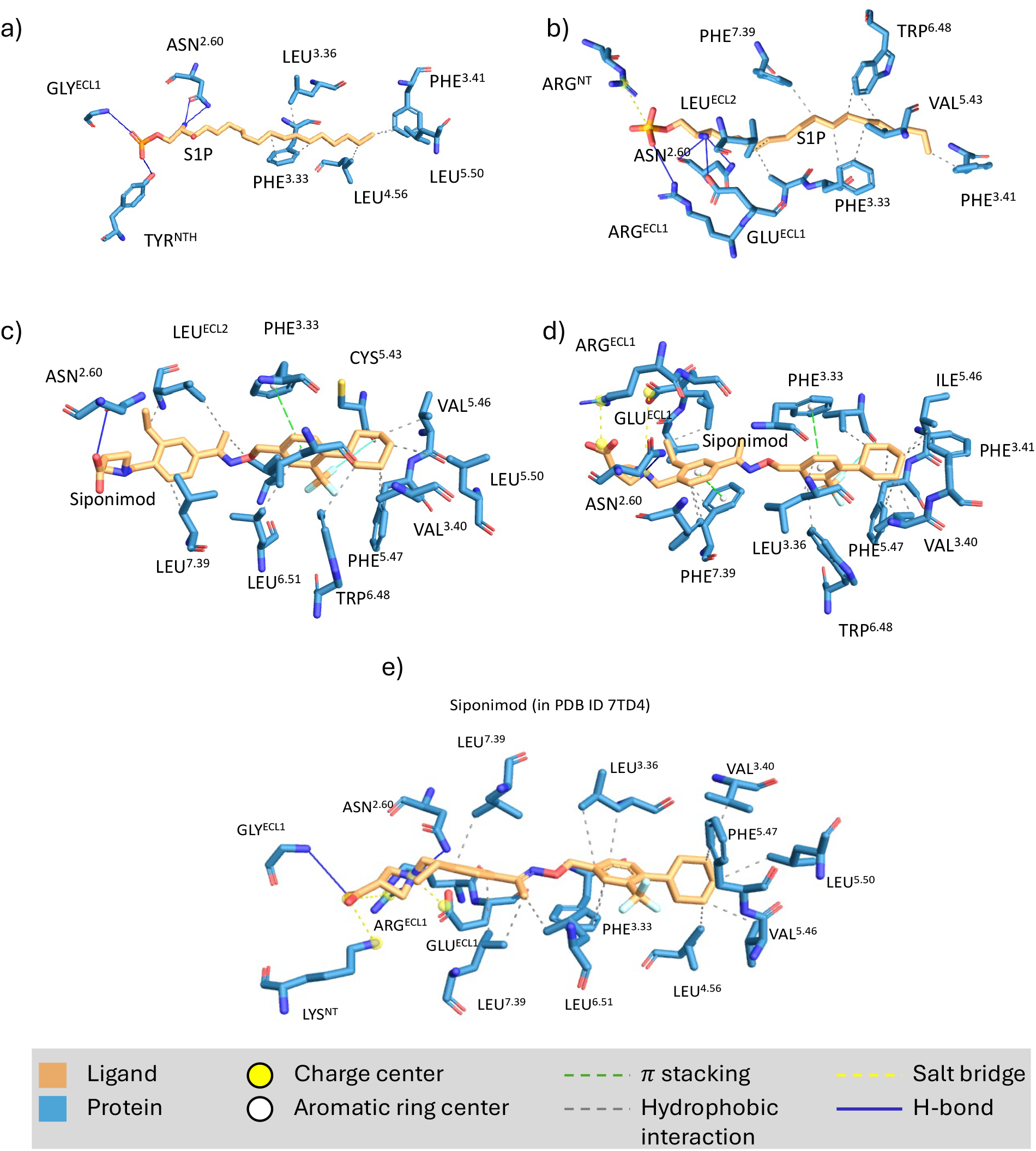
PLIP analysis on the ligand bound S1PR1 and S1PR2 conformations obtained from docking: Hydrophilic head in S1P and Siponimod is engaged in hydrogen bonding with extracellular residues and the aliphatic chain is involved in hydrophobic interactions a) S1P bound S1PR1; b) S1P bound S1PR2; c) Siponimod bound S1PR1; d) Siponimod bound S1PR2; e) Siponimod bound S1PR1 in experimental structure.

S1P engages the N-terminal residues by forming hydrogen bonds and salt bridges with its phosphate head group, and the hydrophobic residues of the receptors interact with the lipid-like tail of S1P (Figure 2b & Figure 2d). In S1PR2, a salt bridge between ARG^*N T*^ and the phosphate head of S1P is also observed. A comparison of the ligand-protein interaction patterns of Siponimod with S1PR1 and S1PR2 shows that Siponimod forms hydrogen bonding interactions with ASN^2.60^ in both S1PR1 and S1PR2. Other than that, Siponimod’s interactions with both the receptors are primarily through weak *π*-stacking and hydrophobic interactions, with additional strong salt bridge interactions with ARG^*ECL*1^ in S1PR2. This explains its higher binding affinity compared to S1PR1 (Figure 2a & Figure 2c).

To validate the docking protocol, the docked conformation of Siponimod in S1PR1 was compared with its conformation in the cryo-EM structure of Siponimod-bound S1PR1 (PDB ID 7TD4). Typically, an RMSD of less than 2 Å between the docked pose and the reference structure (i.e., ligand pose from the crystal structure) is considered sufficiently accurate for ligand docking. ^64,65^ For Siponimod, the hydrophobic tail of the docked pose aligned well with the crystal structure pose, though there was a noticeable shift in the hydrophilic head, resulting in an RMSD (symmetry corrected) of approximately 1.8 Å between the two poses (Figure SI 3), which is acceptable. Since the same docking protocol was used to obtain the Siponimod bound S1PR2 structure, its reliability is supported. However, a comparison of the interactions of docked Siponimod in S1PR1 with experimentally determined Siponimod bound S1PR1 interactions revealed that, due to a different orientation of the polar head group at the N-terminal in the docked structure, interactions with LYS^*N T*^, ARG^*ECL*1^, and GLY^*ECL*1^ were absent in the docked pose of Siponimod in S1PR1 (Figure 2e).

Although a recent study^27^ reveals that the molecular docking analysis of diverse S1PR modulators to the active sites of both S1PR1 and S1PR5 exhibited a good correlation with experimental data, such analyses can offer only a limited insight. The degrees of freedom of the protein residues were not considered during the rigid docking process, resulting in a static depiction of protein-ligand interactions. To understand these interactions more precisely, it is essential to consider the dynamics of the protein upon ligand binding. Therefore, all-atom unbiased MD simulations of both ligand-free and ligand-bound (S1P & Siponimod) S1PR1 and S1PR2 proteins embedded within lipid bilayers were conducted. This approach enables the comparison of (a) the binding free energy of the ligands, (b) the persistence of various protein-ligand interactions, (c) the impact of ligand binding on the structural dynamics of protein residues, and (d) the coordinated motions of the transmembrane helices.

### MD simulations confirm thermodynamic and kinetic favorability of Siponimod binding to the active site of S1PR2

The MD trajectories (two ≈1*µ*s replicas) were examined for the extent of fluctuations. As expected, residues corresponding to the loop regions have higher freedom to move thus showing higher RMSF values in both S1PR1 and S1PR2 systems. This holds true for both replicas (Figures SI 4a-4d). In the case of S1PR1, in the first replica, the RMSD fluctuations stabilized after ≈200 ns in apo and S1P bound receptor and after ≈500 ns for the Siponimod bound receptor (Figure SI 5a & SI 5b). RMSD fluctuations for all the three systems related to S1PR2 stabilized after ≈300 ns (Figure SI 5a & SI 5b). In the second replica, all the systems bound to the ligand reach stability after ≈100 ns (Figure SI 5c & SI 5d). The distribution of the first two principal components (PC1 & PC2), calculated from the entire ≈2*µ*s trajectory combining both replicas and considering all the heavy atoms, indicates that the two replicas explore distinct, non-overlapping regions of the conformational space, specially for the S1PR2 systems (Figure SI 6). To access the convergence of the MD trajectories, we performed Markov State Model (MSM) analysis using PC1 & PC2 and found that the implied timescales plateau with increasing lag time, indicating reliable sampling suitable for studying ligand-induced structural changes in the receptors (Figure SI 8). This is further supported by the Chapman-Kolmogorov (CK) test, which shows acceptable agreement between predicted and observed transition probabilities, confirming the kinetic consistency of the MSM models.

To maintain uniformity, MMGBSA calculations, along with pair-wise residue decomposition analysis, were performed on the last 500 ns (of the total ≈1 *µ*s) trajectory, during which stable RMSD fluctuations were observed in all six cases in both the replicas. The relative binding free energies obtained from MMGBSA method 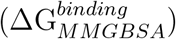 were also obtained for 5 short ≈20 ns replicas and are in good agreement with the values obtained from the longer trajectories (Table 1). 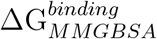 values suggest that Siponimod binds strongly at the active sites of both the receptors S1PR1 and S1PR2, as also suggested by the docking studies. However, the docking studies were unable to capture that the binding strength of Siponimod towards S1PR1 and S1PR2 are comparable. In both the cases, the 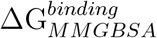 values vary between -60 to -70 kcal/mol.

**Table 1:** Comparison of the ligand binding free energies 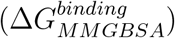 obtained from MMG-BSA calculations performed on different replica of independent MD simulations: entire trajectories of five shorter (≈ 20 ns) replicas and the last 500 ns trajectories of the two longer (≈ 1 *µ*s) replicas. The 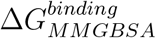 are reported in kcal/mol. The experimentally determined pEC_50_ values are obtained from the PHAROS database.

| $\Delta G_{MMGBSA}^{binding}$ of | Experimental<br>(pEC<sub>50</sub>) | short ($\approx 20$ ns) replicas | | | | | long ($\approx 1 \mu s$) replicas | | Average<br>(kcal/mol) | Docking score<br>(kcal/mol) |
| --- | --- | --- | --- | --- | --- | --- | --- | --- | --- | --- |
|  |  | 1 | 2 | 3 | 4 | 5 | 1 | 2 |  |  |
| S1P to S1PR1 | 10.57 | -94.5 ± 7.6 | -94.9 ± 8.6 | -94.7 ± 6.1 | -91.57 ± 4.8 | -92.7 ± 7.2 | -77.8 ± 5.0 | -74.7 ± 6.4 | -88.7 | -5.8 |
| Siponimod to S1PR1 | 9.41 | -61.8 ± 2.7 | -60.9 ± 2.5 | -61.9 ± 3.0 | -66.8 ± 3.1 | -65.9 ± 3.4 | -60.6 ± 1.0 | -63.4 ± 3.4 | -63 | -11.7 |
| S1P to S1PR2 | 9.64 | -74.1 ± 4.4 | -70.7 ± 5.8 | -69.9 ± 4.4 | -72.7 ± 4.6 | -68.5 ± 4.2 | -68.5 ± 5.0 | -82.7 ± 7.0 | -72.4 | -7.7 |
| Siponimod to S1PR2 | NA* | -67.8 ± 2.5 | -65.9 ± 2.9 | -68.7 ± 2.6 | -68.2 ± 3.3 | -67.1 ± 3.0 | -67.8 ± 2.0 | -60.7 ± 2.9 | -66.6 | -13.0 |

As per the 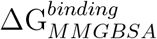 values, the native ligand S1P binds to both receptors with a stronger affinity compared to the approved drug Siponimod (Table 1). Again, binding strength of S1P towards S1PR1 (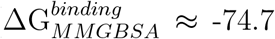 to -94.9 kcal/mol) is higher than that towards S1PR2 (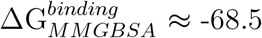 to -82.7 kcal/mol).

Our results also align well with the experimentally reported activity values. The pEC_50_ values for S1P and Siponimod against S1PR1 are 10.57 and 9.41, respectively (source: PHAROS database^66–68^). This indicates a higher affinity of S1P for S1PR1 compared to Siponimod. Additionally, the affinity of S1P for S1PR2 (pEC_50_ = 9.64) is reported to be^22,27^ lower than its affinity for S1PR1 (pEC_50_ = 10.57). However, for Siponimod-S1PR2, the EC_50_ value is reported to be > 10000 nM. As previously mentioned, these values are derived from the GTP*γ*[^35^S] functional assay, which does not necessarily evaluate whether Siponimod can bind to S1PR2, but only evaluates the activation of S1PR2 by Siponimod. Note that, evidence of S1P and Siponimod binding to the same active site in S1PR1 has been reported using competitive radioligand binding assays. ^22,27^ Cryo-EM structure of S1P-bound S1PR2 has also been solved.^17^ But, to our knowledge, there is no reported study on Siponimod binding to S1PR2 using competitive radioligand binding assays or equivalent experimental methods. Hence, the experimental reports remain ambiguous regarding Siponimod’s binding to S1PR2.

MMGBSA calculations mentioned above suggest that Siponimod exhibits favorable binding free energies toward the active sites of both S1PR1 and S1PR2, indicating thermodynamic stability of the ligand-protein complexes. However, MMGBSA primarily captures equilibrium thermodynamic effects and does not provide insights into the kinetic stability of the bound ligand. In other words, while a ligand may bind favorably from a free energy standpoint, it could still exhibit a short residence time if the binding is not kinetically stable.^69,70^ Since our simulations begin from pre-formed ligand-bound states, it is critical to assess whether the ligand remains stably bound over time or tends to dissociate readily.

To address this, we performed *τ*RAMD (Random Acceleration Molecular Dynamics) simulations, which are well-suited for estimating relative residence times of ligands within binding pockets. The *τ*RAMD method introduces an external random acceleration to expedite ligand unbinding events, thereby enabling comparative evaluation of kinetic stability across systems within accessible simulation timescales. The resulting residence times, averaged over five independent replicas, are presented in Table SI1 & Figure SI 9. The significantly longer residence time of Siponimod in S1PR2 (18.0 ns) compared to S1PR1 (8.5 ns) suggests that, in addition to being thermodynamically favorable, the Siponimod–S1PR2 complex is also kinetically more stable. These results further substantiate that Siponimod binds in the S1PR2 active site, as observed in our docking and MMGBSA analyses as well.

Computational studies performed in this work suggest that Siponimod can bind to the active site of S1PR2, while experimental reports indicate that Siponimod cannot modulate S1PR2 activity. To reconcile these findings, we aim to investigate whether Siponimod indeed binds to the active site of S1PR2 but fails to induce the structural changes necessary for modulation. The following sections will delve into these investigations, exploring the underlying mechanisms and potential implications.

### Siponimod utilizes different binding strategies at the active sites of S1PR1 and S1PR2

We found that during the Siponimod–S1PR1 system simulations, the polar head group of Siponimod remains well-anchored, whereas its hydrophobic tail adopts a slightly shifted orientation relative to the crystal structure, likely due to strong local interactions within the hydrophobic pocket that restrict its flexibility (Figure SI 3g & SI 3h). Nevertheless, Siponimod and S1P appear to employ a similar overall strategy for engaging the active site of S1PRs. The polar head region of S1P and Siponimod engages in electrostatic interactions with active site residues, while the nonpolar tail is stabilized by hydrophobic interactions. The phosphate group on the S1P ligand forms salt bridges and strong hydrogen bonds with the N-terminal region of the receptor. Similarly, Siponimod’s polar head is stabilized within the pocket through hydrogen bonding interactions. However, a more detailed investigation reveals notable differences between the two ligands’ binding modes.

Pairwise energy decomposition analysis (Figure 3a & 3b) reveals that in S1PR1, there are 15 common residues that significantly contribute to stabilizing both S1P and Siponimod binding at the active site. Additionally, four specific residues (SER105, TYR29, ASN101, and ARG120) interact strongly with S1P but do not engage with Siponimod. Conversely, three residues (MET124, ALA293, and GLU294) interact exclusively with Siponimod.

**Figure 3:**
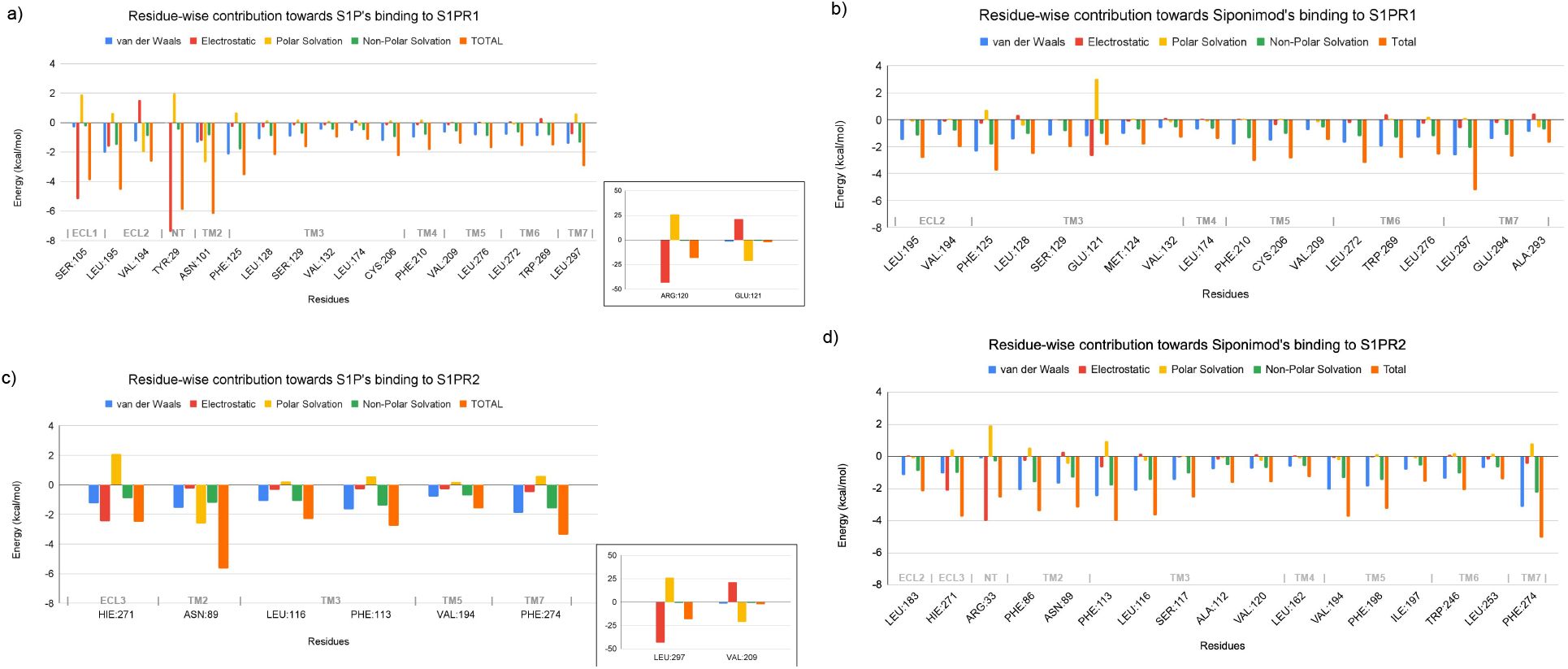
Residue-wise contribution towards binding free energies of (a) S1P to S1PR1, (b) Siponimod to S1PR1, (c) S1P to S1PR1 and (d) Siponimod to S1PR2, averaged over 2 independent replicas of long (≈1*µ*s each) MD trajectories. Most of the energy components fall within the range of -8 to 4 kcal/mol. Instances where one or more energy components exceed this range are displayed separately in the inset.

In the case of S1PR2, S1P interacts strongly with only 8 residues, of which ARG108 uniquely binds to S1P (Figure 3a & 3b). The remaining 7 residues also participate in stabilizing Siponimod at the active site of S1PR2. Moreover, Siponimod engages with an additional 10 residues in S1PR2’s active site, including LEU183, PHE86, SER117, ALA112, VAL120, LEU162, PHE198, ILE197, TRP246, and LEU253.

Although the similar set of active site residues contribute towards binding of S1P and Siponimod, the chemical nature of interaction varies between the binding of S1P and Siponimod. While S1P binding to the active site is primarily driven by electrostatic interactions, van der Waals forces play a key role in stabilizing Siponimod. Approximately 55% of Siponimod’s binding free energy to S1PR1 (and about 52% to S1PR2) arises from a network of individually weak van der Waals interactions with residues across various secondary structural domains. The most significant contributions come from LEU297 in S1PR1 (L^7.39^) and PHE274 in S1PR2 (F^7.39^), a residue previously believed to hinder Siponimod’s binding to the active site of S1PR2. In contrast, a few strong electrostatic interactions (such as ARG120 in S1PR1 and ARG108 in S1PR2) are responsible for anchoring S1P at the active site.

Analysis of pairwise energy decomposition (Figure 3) further suggests that Siponimod’s binding profiles to S1PR1 and S1PR2 differ significantly. In S1PR1, nearly 40% of Siponimod’s binding free energy stems from its interactions with residues in TM6 and TM7, whereas in S1PR2, this contribution is only about 18%. Among the TM6 and TM7 residues that interact exclusively with Siponimod in S1PR1, LEU272 (L^6.51^) contributes the most (−3.2 kcal/mol). The TM6/TM7 residues shared between S1PR1 and S1PR2 include W^6.48^ (part of a key transmission switch, discussed later) and L^7.39^, which is mutated to F^7.39^ in S1PR2. These subtle differences likely contribute to limiting Siponimod’s potential as a modulator of S1PR2.

### The binding of Siponimod and S1P differentially influences the structural dynamics of S1PR2

Dynamic cross-correlation maps (DCCM) provide an averaged representation of how each residue moves relative to others throughout the simulation. DCCMs were generated for the HOLO systems taking the entire ≈ 2*µ*s data combining both the replicas (Figure SI 10). Both S1P (Figure SI 10a) and Siponimod (Figure SI 10b) binding to S1PR1 induce similar patterns of correlated and anti-correlated motions across the receptor residues. The only notable difference occurs in the dynamics of the N-terminal (NT) domain. Upon Siponimod binding, the NT residues exhibit correlated movements with residues from the extracellular loops ECL1 and ECL2 (region encircled in orange and brown in Figure SI 10b), whereas these movements are anti-correlated when S1P is bound. However, the DCCMs show two major differences in mutual structural dynamics of S1PR2 residues that is induced by Siponimod binding exclusively. They are discussed in the following sections.

The first difference is observed in the dynamics of the intracellular loops ICL2 and ICL3 with respect to the rest of the protein. ICL3 is the longest cytoplasmic loop in S1PR1, whereas in S1PR2, both ICL2 and ICL3 are of similar length. When S1P binds, the residues of ICL3 (and part of TM6) in S1PR1 (Figure SI 10a, region encircled in grey) and the residues of ICL2 in S1PR2 (Figure SI 10c, region encircled in green) exhibit strong anti-correlated movement with the rest of the receptor. Siponimod binding replicates this anti-correlated dynamic in S1PR1 (Figure SI 10b, region encircled in grey), but fails to induce a comparable effect in S1PR2 (Figure SI 10d, region encircled in green). Note that, ICL2 and ICL3 in S1PRs interact with the G*α* protein (not part of the MD systems) in the cytoplasm.^71,72^ Hence, although Siponimod binds tightly to the active site, it does not significantly influence the dynamics of the G protein-binding regions in S1PR2.

The second difference is found in the mutual dynamics of transmembrane (TM) helices. When S1P binds to S1PR1 or S1PR2, the TM helices display mostly anti-correlated motions. However, when Siponimod binds to S1PR2, several motion patterns shift, leading to moderately correlated motions, such as between TM6 and TM2 (Figure SI 10d, region encircled in black). The TM2 also shows correlated motions with ICL1 on Siponimod binding (Figure SI 10d, region encircled in magenta). Siponimod binding to S1PR1 does not cause similar changes. These findings highlight the differential impact of Siponimod and S1P binding on S1PR2’s dynamic behavior.

### Key transmission switches remain inactive upon Siponimod binding to S1PR2

Activation of GPCRs, including the five S1P receptor family members, is known to take place through conformational changes of certain residues in the transmembrane helices. These residues are generally referred to as transmission switches. ^30,73–75^ Through MD simulations, we sought to determine the movements of these transmission switches across apo, S1P-bound and Siponimod-bound receptors, S1PR1 and S1PR2. Analysis of MD trajectories reveal that, compared to S1P, Siponimod leads to a different conformation change in the transmission switches upon binding to S1PR2. The differential influence of S1P and Siponimod binding to the two key transmission switches in S1PRs known to differentiate the active vs inactive conformations are discussed below.

1. W^6.48^ serves as a critical switch in GPCRs, including the S1P receptors, leading to receptor activation upon ligand binding.^76,77^ Careful observations of the MD trajectories reveal that the W^6.48^ switch is present in two distinctly different orientations when it is bound to S1P (active mode) vs. when ligand free (inactive mode). In S1PR1, the W^6.48^ switch is found to stay in its active mode during the trajectory when Siponimod is bound (Figure 4a). However, in S1PR2, the W^6.48^ switch is no longer found in its active mode once Siponimod is bound (Figure 4b). This change in W^6.48^ switch conformation in Siponimod bound S1PR2 resulted in an inward movement of TM6 resulting in a decrease in distance between TM3 and TM6 as evident from the distance between V^3.40^ and F^6.44^ (Figure 5).
2. Another conserved motif, P^5.50^-I^3.40^-F^6.44^, in GPCRs is known to play important role in receptor activation.^78,79^ In case of S1PR1 and S1PR2, L^5.50^-V^3.40^-F^6.44^ and I^5.50^-V^3.40^-F^6.44^ motifs, respectively, play a similar role.^80^ Similar to the W^6.48^ switch, the L^5.50^- V^3.40^-F^6.44^ switch in S1PR1 and the I^5.50^-V^3.40^-F^6.44^ switch in S1PR2, are present in two distinctly different orientations when they are bound to S1P (active mode) vs. when ligand free (inactive mode). In S1PR1, the L^5.50^-V^3.40^-F^6.44^ switch maintains its active mode (marked by F^6.44^ and V^3.40^ orientations) when Siponimod is bound (Figure 4c). However, in S1PR2, the I^5.50^-V^3.40^-F^6.44^ switch could not retain its active mode (marked by a shift in F^6.44^ orientations) on Siponimod binding (Figure 4d).

**Figure 4:**
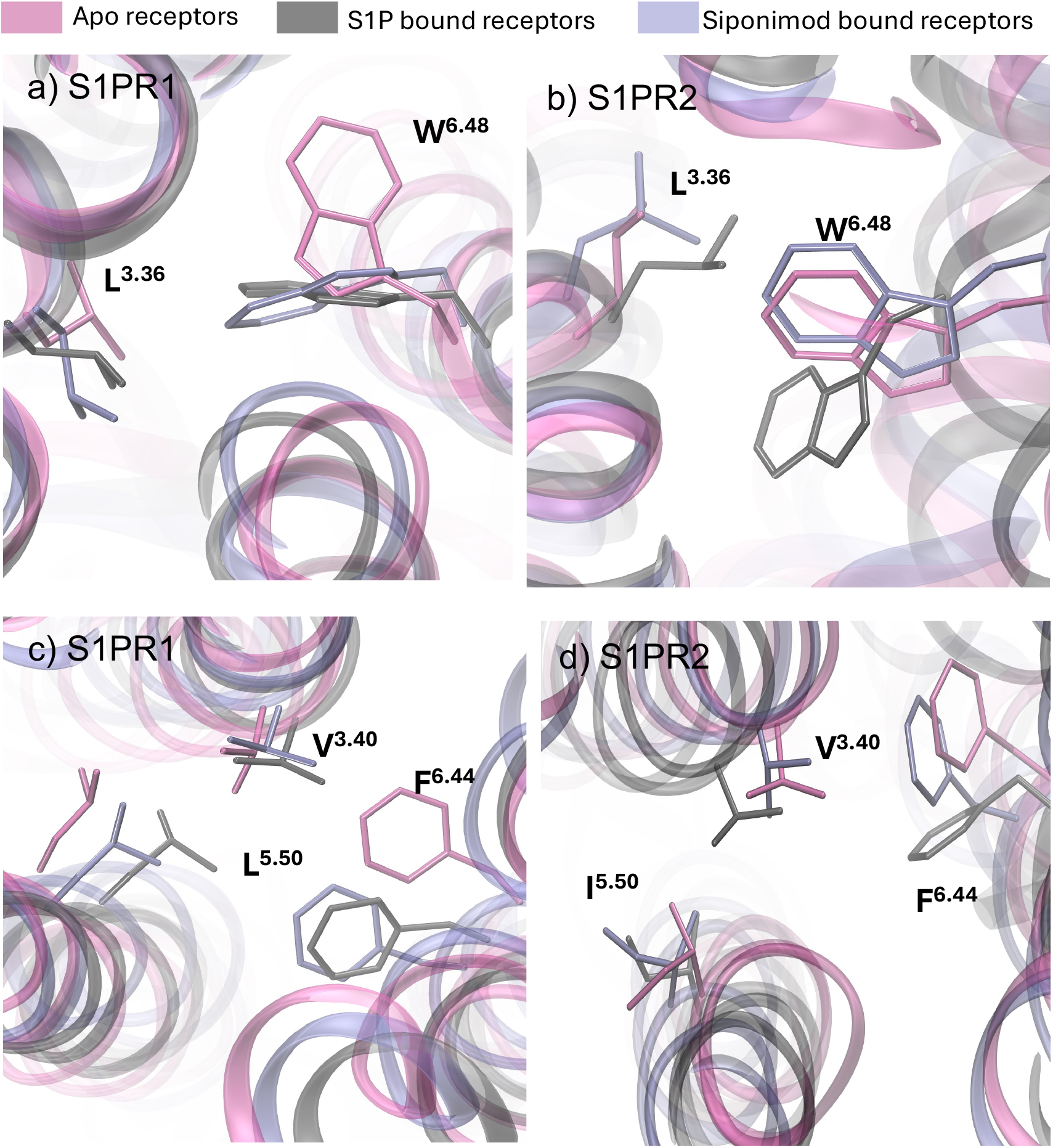
Transmission switches in S1PR1 and S1PR2. a) With respect to the respective APO systems L^3.36^-W^6.48^ switches exhibit similar flip in S1P and Siponimod bound S1PR1; L^3.36^-W^6.48^ switches exhibit different orientations in S1P and Siponimod bound S1PR2 (the orientation in Siponimod bound system resembles the APO system) ; c) With respect to the APO system, S1P and Siponimod exhibiting similar structural changes in L^5.50^-V^3.40^-F^6.44^ motif in S1PR1; d) S1P and Siponimod exhibiting noticeably different structural changes in I^5.50^-V^3.40^-F^6.44^ motif (the orientation in Siponimod bound system resembles the APO system). The last 100 frames from replica 1 of each system were analyzed to select the most illustrative conformational snapshots. Customized collective variables (CVs) were employed to track the switching events described above across the MD trajectories of both replicas (see Figures SI 11 and SI 12)

**Figure 5:**
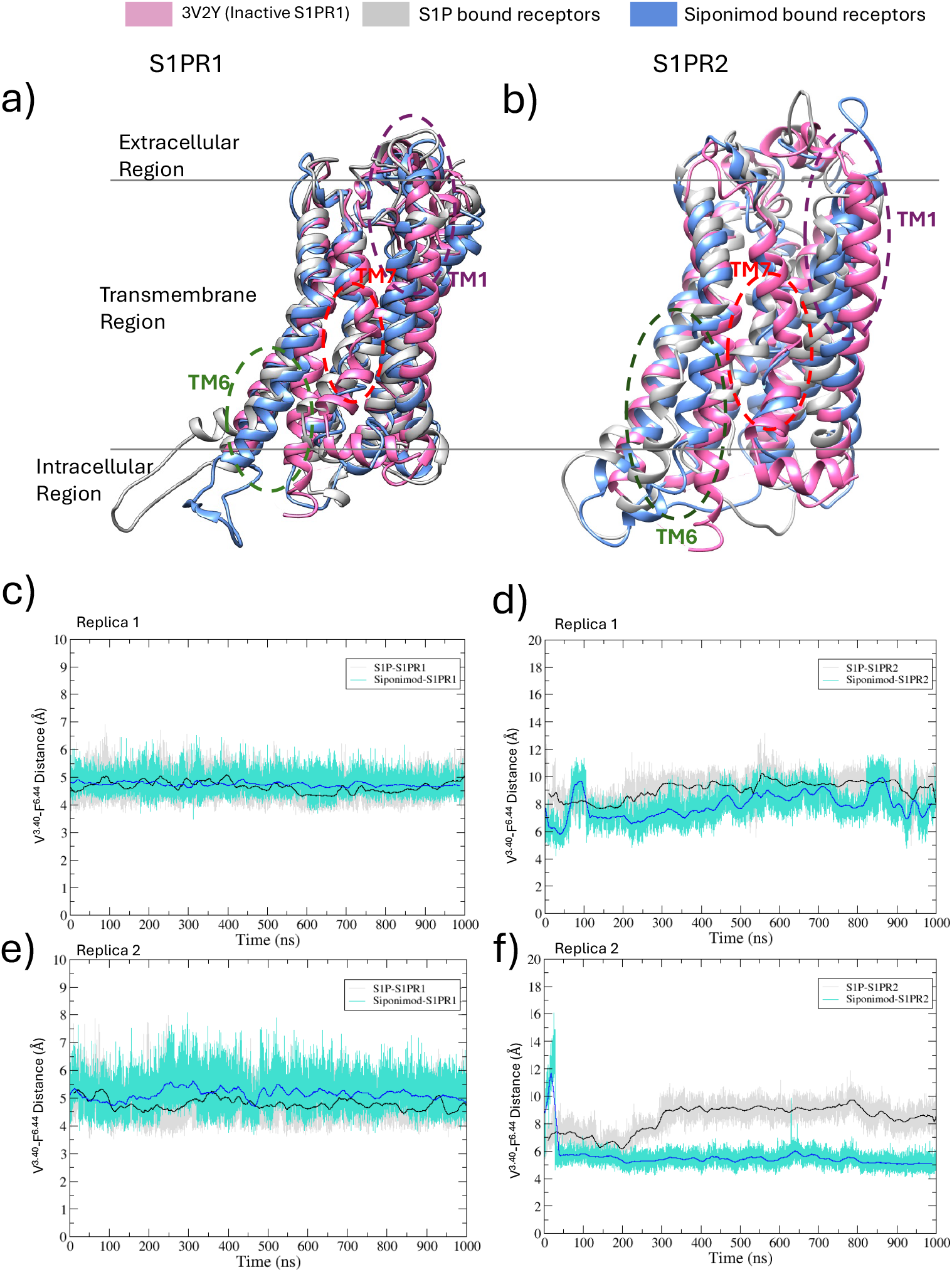
In panels (a) and (b), we compare the S1P- and Siponimod-bound conformations of S1PR1 and S1PR2 with the inactive S1PR1 structure. Panels (c) and (e) show the distance changes between helices TM3 and TM6 for S1PR1 across both replicas, while panels (d) and (f) show the same for S1PR2. In S1PR1, TM6 remains mostly outward in both the S1P and Siponimod-bound states. However, in Siponimod-bound S1PR2, the distance between TM3 and TM6 decreases due to TM6 moving inward. TM6 tends to move inward in Siponimodbound S1PR2 but stays outward in S1P-bound S1PR2. The distance between TM3 and TM6 remains consistent in both S1P-bound S1PR1 and S1PR2, as well as in Siponimod-bound S1PR1.

### Siponimod binding drives TM6–TM7 rearrangement away from the active-state in S1PR2

The relative movements of TM6 and TM7 in S1P receptors are closely linked to receptor activation.^17,81^ Our analysis shows that ligand binding triggers significant conformational changes in key residues, particularly F^6.44^ and V^3.40^. These shifts, coupled with movements in W^6.48^ and L^3.36^, play a crucial role in regulating the relative positioning of the transmembrane helices, which ultimately influences receptor activation dynamics. The DCCM plots (Figure SI 10) reveal that TM6 residues exhibit distinct motion in S1PR2 when Siponimod is bound, in contrast to the S1P-bound state. Porcupine plots obtained from PCA of the MD trajectories (Figure SI 7) suggests that in response to S1P binding at the active site, TM6 and TM7 move away from each other in opposite directions in both S1PR1 and S1PR2. On Siponimod binding, similar movements of TM6 and TM7 residues are observed in S1PR1, but not in S1PR2.

So, a similar transmission switch dynamics is observed when Siponimod binds to S1PR1, but not when it binds to S1PR2 as compared to their respective S1P-bound counterparts. Given that the MD simulations were initiated from an active state conformation of the receptors, this implies that Siponimod binding at the active site of S1PR2 is not able to hold the active state conformation of S1PR2. The relative positioning of key transmembrane helices, specifically TM6 and TM7, was analyzed in relation to the known inactive conformation of S1PR1 (PDB ID: 3V2Y^82^). The last frames from the two MD simulation replicas were over-laid onto the inactive S1PR1 conformation (Figure 5). It is evident that S1PR2 resembles the inactive state conformation (pink structure in Figure 5b) when it is bound to Siponimod (blue structure in Figure 5b). S1P bound S1PR2 conformation is distinctly different.

To quantify the positional shift of TM6 in response to ligand binding, we monitored the distance between the C*β* atoms of residues V^3.40^ and F^6.44^ (Figure 5). This metric captures the inward movement of TM6 relative to TM3, a hallmark of receptor deactivation. In the Siponimod-bound S1PR2 system, this distance decreases noticeably, particularly in replica 2 (Figure 5)e), suggesting an inward drift of TM6. While the change is less pronounced in replica 1 (Figure 5)d), a subtle inward shift is still observed when compared to the Siponimod-bound S1PR1 system, which remains conformationally stable throughout the simulation (Figure 5a & 5b).

To further validate this trend, we performed an additional 500 ns simulation (replica 3) for both S1P- and Siponimod-bound S1PR2 systems. As shown in Figure SI 14, the new replica confirms the inward movement of TM6 in the Siponimod-bound state, whereas the S1P-bound receptor maintains a stable V^3.40^–F^6.44^ distance, consistent with its active-state conformation.

These observations support the notion that Siponimod binding to S1PR2 promotes a partial transition toward an orientation observed in inactive state, reminiscent of the experimentally determined inactive structure of S1PR1. This structural drift likely results from weaker interactions between Siponimod and the TM6 and TM7 residues in S1PR2 (Figure 3), further contributing to its inability to stabilize the active conformation.

### TM7 residues in S1PR2 adopt ligand-specific conformations that influence selectivity

In addition to the transmission switches mentioned earlier, significant differences in the conformation of residues near the extracellular region of TM7 were observed between S1P-bound and Siponimod-bound S1PR2. These differences are mainly driven by the dynamics of H^7.36^, Y^7.37^, F^7.38^, and F^7.39^ in the TM7 helix. Although the orientation of these residues varies slightly between the two replicas, the relative positioning of H^7.36^ and F^7.39^ remains consistent (Figure SI 13). Notably, these residues are absent in S1PR1, as discussed earlier. It appears that conformational changes in H^7.36^ and the associated residues (Y^7.37^, F^7.38^, and F^7.39^) contribute to S1PR2 activation upon S1P binding, but not with Siponimod binding.

We analyzed the *χ*_1_ vs. *χ*_2_ dihedral angle distributions of H^7.36^, Y^7.37^, F^7.38^, and F^7.39^ in the S1P- and Siponimod-bound S1PR2 systems (Figure 6). All simulations were initiated from the active-state structure of S1P-bound S1PR2, with dihedral values from the experimental model indicated in each panel. Red dashed circles mark the conformational basin corresponding to this active state. Both ligands show some sampling near the starting conformation, but clear differences emerge. S1P-bound S1PR2 explores additional regions (orange dashed circles) not sampled in the Siponimod-bound system, while Siponimod leads to unique conformations absent in the S1P-bound state (blue dashed circles). These patterns indicate that the key TM7 residues adopt distinct conformational preferences depending on the ligand. S1P promotes side-chain orientations associated with activation, whereas Siponimod favors alternative states, supporting the view that it fails to trigger the same transmission switch rearrangements in S1PR2.

**Figure 6:**
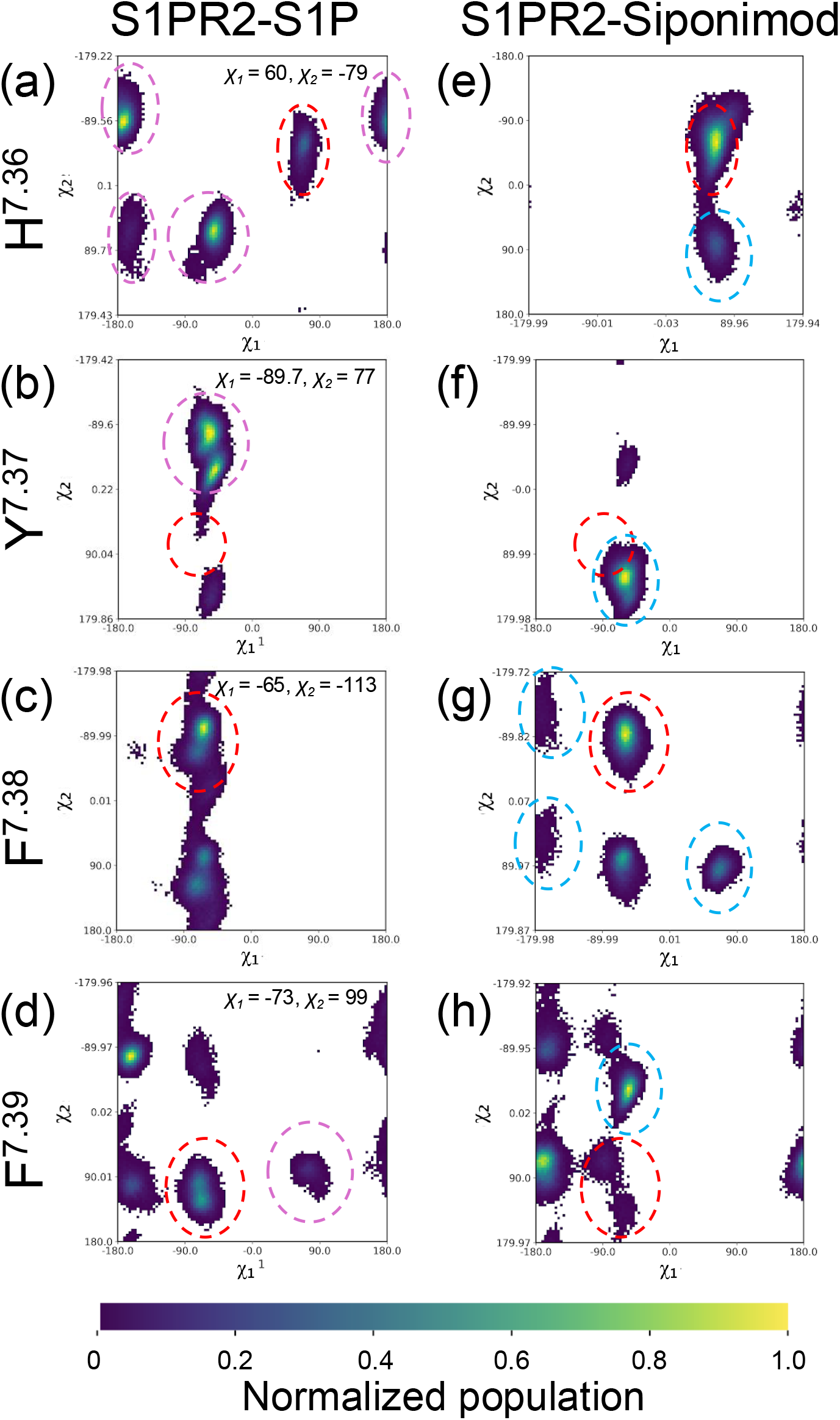
Comparison of side-chain dihedral angles for key differentiating residues in TM7 of S1PR2. The side chains of residues H^7.36^, Y^7.37^, F^7.38^, and F^7.39^ in the TM7 helix adopt distinct orientations in the S1P-bound S1PR2 complex (panels a−d) compared to the Siponimod-bound S1PR2 complex (panels e−h).

Previous studies^17^ have also shown that PHE^7.39^ in S1PR2 plays a role in its selectivity towards the inhibitor JTE-013. Specifically, the mutation F274I reduced the receptor’s response to JTE-013 but enhanced downstream activity upon binding to FTY720-P. Thus, we hypothesize that PHE^7.39^, along with HIS^7.36^ and TYR^7.37^, could be critical in designing selective modulators for S1P receptors. Another key residue in this context is L^6.51^, which makes the largest contribution to Siponimod’s binding to S1PR1 but does not interact with Siponimod in S1PR2. These residues could be targeted to optimize molecules for either enhanced or reduced binding to S1PR2, depending on therapeutic requirements.

## Conclusion

Our simulations reveal that Siponimod exhibits strong binding affinity and prolonged residence time at the active site of S1PR2, as demonstrated by MMGBSA calculations and enhanced *τ* -RAMD simulations. However, despite this apparent thermodynamic and kinetic stability, Siponimod-bound S1PR2 fails to exhibit the conformational rearrangements typically associated with receptor activation. In particular, key transmission switches—W^6.48^ and the I^5.50^–V^3.40^–F^6.44^ triad—remain inactive, and the outward movement of TM6 and TM7, observed in S1P-bound systems, is absent. These findings lead us to hypothesize that although Siponimod binds to S1PR2, it is unable to stabilize the active-state conformation required for functional signaling. This decoupling of binding from activation likely underlies its functional selectivity and lack of potential to modulate S1PR2.

To experimentally validate this hypothesis, we recommend performing competitive radioligand binding assays to directly assess Siponimod’s ability to bind to S1PR2. Similar assays have previously confirmed Siponimod’s binding to S1PR1 and S1PR5,^27^ and extending this approach to S1PR2 would provide essential experimental evidence to support or refute our computational observations.

Our findings underscore the need to incorporate structural and dynamic signatures—beyond static binding energies—when evaluating drug candidates using *in silico* approaches. In particular, features such as transmission switch activation, TM6–TM7 rearrangements, and side-chain flexibility in residues like PHE^7.39^, HIS^7.36^, and TYR^7.37^ can serve as critical determinants of receptor-specific ligand response. These residues could be exploited in future efforts to design selective modulators for S1P receptors.

Nonetheless, our work has limitations. The absence of G-protein partners in the simulation systems may impact the long-term stability of the active state, as reported for other GPCRs like the *β*_2_-adrenergic receptor.^83,84^ Capturing the full activation mechanism or conformational equilibrium between active and inactive states would require longer simulations, more replicas, or enhanced sampling techniques—all of which are avenues for future work. Despite these limitations, our study demonstrates how atomistic simulations can offer mechanistic insights into GPCR selectivity and support structure-based drug design for challenging targets.

## Acknowledgement

We appreciate the contribution of Toran Roy and Srinivas Tumuluri in the initial discussions on the peoject. The authors thank Dr. Shubhandra Tripathi, Dr. Madhura Mohole, Dr. Srinath V. K. Kompella, Prasad Chodavarapu and Vikram Duvvoori for the invaluable discussions and constructive feedback during the preparation of manuscript. We sincerely thank the reviewers for their insightful comments and constructive suggestions, which greatly helped improve the clarity and rigor of this manuscript.

## Data and Software Availability

Molecular docking studies were conducted using the AutoDock VINA tool (v1.1.2) within UCSF Chimera (v1.16.0). Molecular dynamics (MD) simulations were performed with GROMACS (v2022.3). To ensure reproducibility, all input files, parameter files, and scripts used for system preparation and performing MD simulations are shared as supporting information (JCIM-publication-data.zip).

## Supporting Information Available

See the “Supporting Information” material includes Figures SI 1–SI 14 and Table SI 1, along with details of the MD protocol, sequence and structure alignments, ligand docking, RMSD/RMSF and PCA analyses, Markov state modeling, *τ* -RAMD residence time calculations, residue-wise binding energy decomposition, inter-residue correlation maps, transmission switch analyses across replicas, and a list of collective variables used in this study.

## TOC Graphic

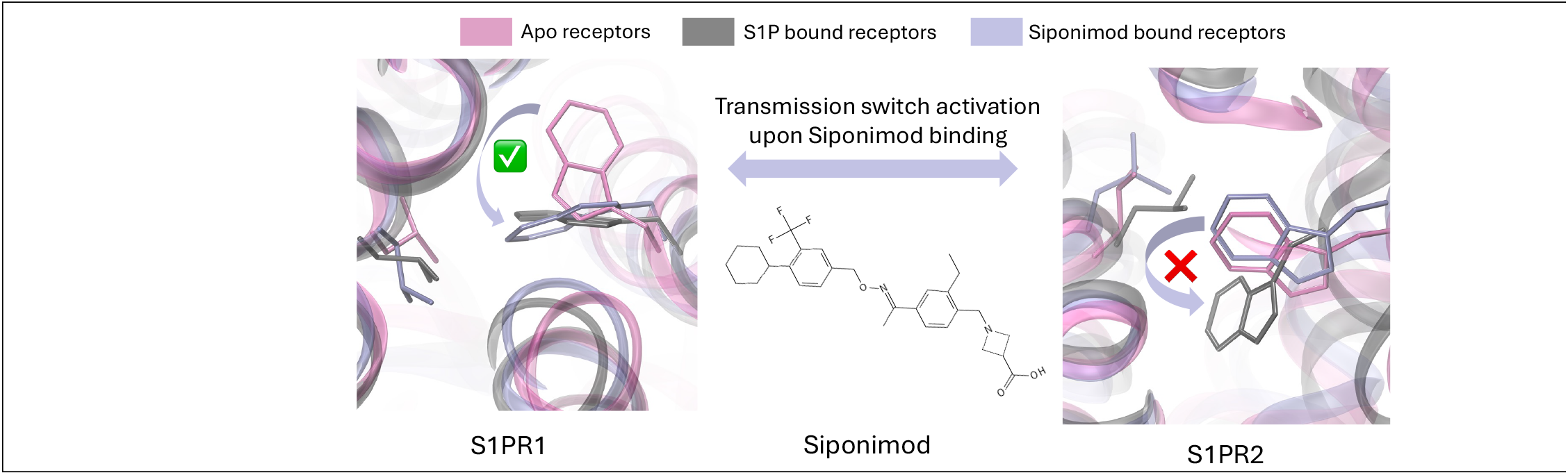

