## Supporting Information for "Selective Activation of GPCRs: Molecular Dynamics Shows Siponimod Binds but Fails to Activate S1PR2 Unlike S1PR1"

### 1. Details of Molecular Dynamics (MD) Protocol used

The MD systems were prepared and minimized using AmberTools22. Amber compatible files were converted to GROMACS compatible files and the systems were further minimized in GROMACS 2022.3. For all MD simulations conducted in this study, the steepest descent algorithm was employed for energy minimization. A maximum force threshold of less than 1000 kJ/mol/nm was set as the convergence criterion. NVT equilibration for 1 ns was carried out using velocity rescaling thermostat[1]. NPT equilibration for 1 ns followed by production run in NPT were carried out using Parrinello-Rahman barostat[2], [3]. The Particle Mesh Ewald (PME) algorithm[4] was used to account for the long-range electrostatic interactions. Leap-frog algorithm with a time step of 2 fs was used for MD integration. The temperature and pressure were maintained at 300 K and 1 bar, respectively. Hydrogen atoms were constrained using the LINCS algorithm[5].

### 2. Sequence Alignment of S1PR1 with Respect to S1PR2 and the Residues Important for Their Activation Mechanism

| Score | Expect | Method | Identities | Positives | Gaps |
|---|---|---|---|---|---|
| 328 bits(841) | 1e-115 | Compositional matrix adjust. | 162/323(50%) | 223/323(69%) | 13/323(4%) |

```
Query  17   SDYVNYDIIVRHYNYTGKLNISADKENSIKLTSVVFILICCFIILENIFVLLTIWKTKKF  76
            S+Y+N + +  HYNYT K  +   +  S ++ S   +++CC I++EN+ VL+ + +  KF
Sbjct  6    SEYLNPNKVQEHYNYT-KETLETQETTSRQVASAFIVILCCAIVVENLLVLIAVARNSKF  64

Query  77   HRPMYYFIGNLALSDLLAGVAYTANLLLSGATTYKLTPAQWFLREGSMFVALSASVFSLL  136
            H  MY F+GNLA SDLLAGVA+ AN LLSG+ T +LTP QWF REGS F+ LSASVFSLL
Sbjct  65   HSAMYLFLGNLAASDLLAGVAFVANTLLSGSVTLRLTPVQWFAREGSAFITLSASVFSLL  124

Query  137  AIAIERYITMLKMKLHNGSNNFRLFLLISACWVISLILGGLPIMGWNCISALSSCSTVLP  196
            AIAIER++ + K+KL+    + R+ LLI A W+ISL+LGGLPI+GWNC+  L +CSTVLP
Sbjct  125  AIAIERHVAIAKVKLYGSDKSCRMLLLIGASWLISLVLGGLPILGWNCLGHLEACSTVLP  184

Query  197  LYHKHYILFCTTVFTLLLLSIVILYCRIYSLVRTRSRRLTFRKNISKASRSSEKSLALLK  256
            LY KHY+L   T+F+++LL+IV LY RIY +VR+           S A  ++ ++LALLK
Sbjct  185  LYAKHYVLCVVTIFSIILLAIVALYVRIYCVVRS-----------SHADMAAPQTLALLK  233

Query  257  TVIIVLSVFIACWAPLFILLLLDVGCKVKTCDILFRAEYFLVLAVLNSGTNPIIYTLTNK  316
            TV IVL VFI CW P F +LLLD  C V +C IL++A YF  ++ LNS  NP+IYT  ++
Sbjct  234  TVTIVLGVFIVCWLPAFSILLLDYACPVHSCPILYKAHYFFAVSTLNSLLNPVIYTWRSR  293

Query  317  EMRRAFIRIMSCCKCPSGDSAGK  339
            ++RR  +R + C + P     G+
Sbjct  294  DLRREVLRPLQCWR-PGVGVQGR  315
```

Residues involved in. differentiating Siponimod modulation of S1PR1 and S1PR2

**Figure SI 1.** Sequence alignment result for S1PR1 (Query) with S1PR2 (Subject) as obtained from NCBI BLASTp server. Residues highlighted in orange are used for the BW naming convention. Residues highlighted in green are the ERY(H) and NPxxY signature motifs for GPCRs. Residues highlighted in blue depict the important transmission switches. Residues in pink turned out to be of significance for S1PR1 selectivity as discussed in main manuscript.

**Discussion:** Crucial residues, including transmission switches and conserved moieties, are highlighted in the sequence alignment plot. While most of the residues involved with ligand binding and activation mechanism are similar, a few modifications were seen. As per BW numbering scheme[6] the 50$^{th}$ residue in 5$^{th}$ helix is leucine (L) in S1PR1 while the same role is played by isoleucine (I) in S1PR2, as highlighted in orange in Figure SI 1. Similar difference is observed in the ERY(H) motif where aromatic tyrosine (Y) residue in S1PR1 is replaced by another aromatic but slightly basic histidine (H) residue in S1PR2, as highlighted in green in Figure SI 1. Unlike the above two examples where the modified amino acids are of same physicochemical property, in TM7, a glutamate residue (E) with a negatively charged side chain in S1PR1 is replaced by a histidine (H) with a positively charged side chain in S1PR2, as highlighted in pink in Figure SI 1.

### 3. S1PR1 and S1PR2 Structure Alignment using Foldseek

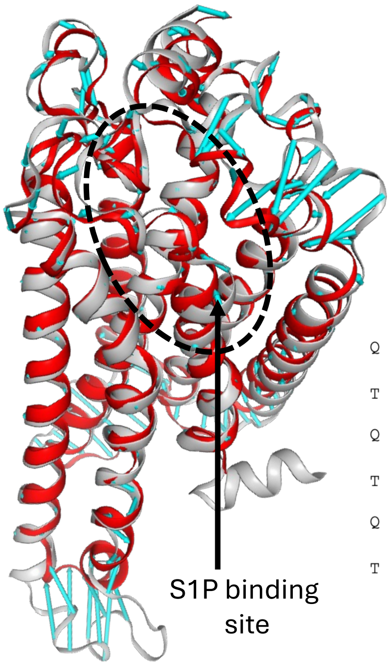

TM-Score: 0.88436

| | Sequence Similarity | Score | E-value | Query position | Target position |
|---|---|---|---|---|---|
| S1PR1-S1PR1 | 1 | 2223 | 1.55e-41 | 1-302 (302) | 1-302 (302) |
| S1PR1-S1PR2 | 0.517 | 830 | 5.46e-19 | 1-290 (302) | 1-278 (279) |

```
Q   1 DIIVRHYNYTGKLNISADKENSIKLTSVVFILICCFIILENIFVLLTIWKTKKFHRPMYYFIGNLALSDLLAGVAYTANL
      + +  HYNYT K  +   +  S ++ S    +++CC I++EN++VL+ + +  KFH  MY F+GNLA SDLLAGVA+ AN
T   1 NKVQEHYNYT-KETLETQETTSRQVASAFIVILCCAIVVENLLVLIAVARNSKFHSAMYLFLGNLAASDLLAGVAFVANT

Q  81 LLSGATTYKLTPAQWFLREGSMFVALSASVWSLLAIAIERYITMLKMKLHNGSNNFRLFLLISACWVISLILGGLPIMGW
      LLSG+ T +LTP QWF REGS F+ LSASV+SLLAIAIER++ + K+KL+    + R++LLI A W+ISL+LGGLPI+GW
T  80 LLSGSVTLRLTPVQWFAREGSAFITLSASVFSLLAIAIERHVAIAKVKLYGSDKSCRMLLLIGASWLISLVLGGLPILGW

Q 161 NCISALSSCSTVLPLYHKHYILFCTTVFTLLLLSIVILYCRIYSLVRTRSRRLTFRKNISKASRSSEKSLALLKTVIIVL
      NC+  L +CSTVLPLY KHY+L   T+F+++LL+IV LY RIY +VR+         +++        +LALLKTV IVL
T 160 NCLGHLEACSTVLPLYAKHYVLCVVTIFSIILLAIVALYVRIYCVVRSS------HADMAAPQ-----TLALLKTVTIVL
```

**Figure SI 2**. Structure comparison of S1PR1 and S1PR2 using Foldseek. S1PR1 (grey) and S1PR2 (red) show a same structural fold as expected of the sub-family. Difference in loop structures in the two cases is shown using blue vectors.

### 4. Aligning Docked Pose of Siponimod to Siponimod in Experimentally Available S1PR1 Receptor

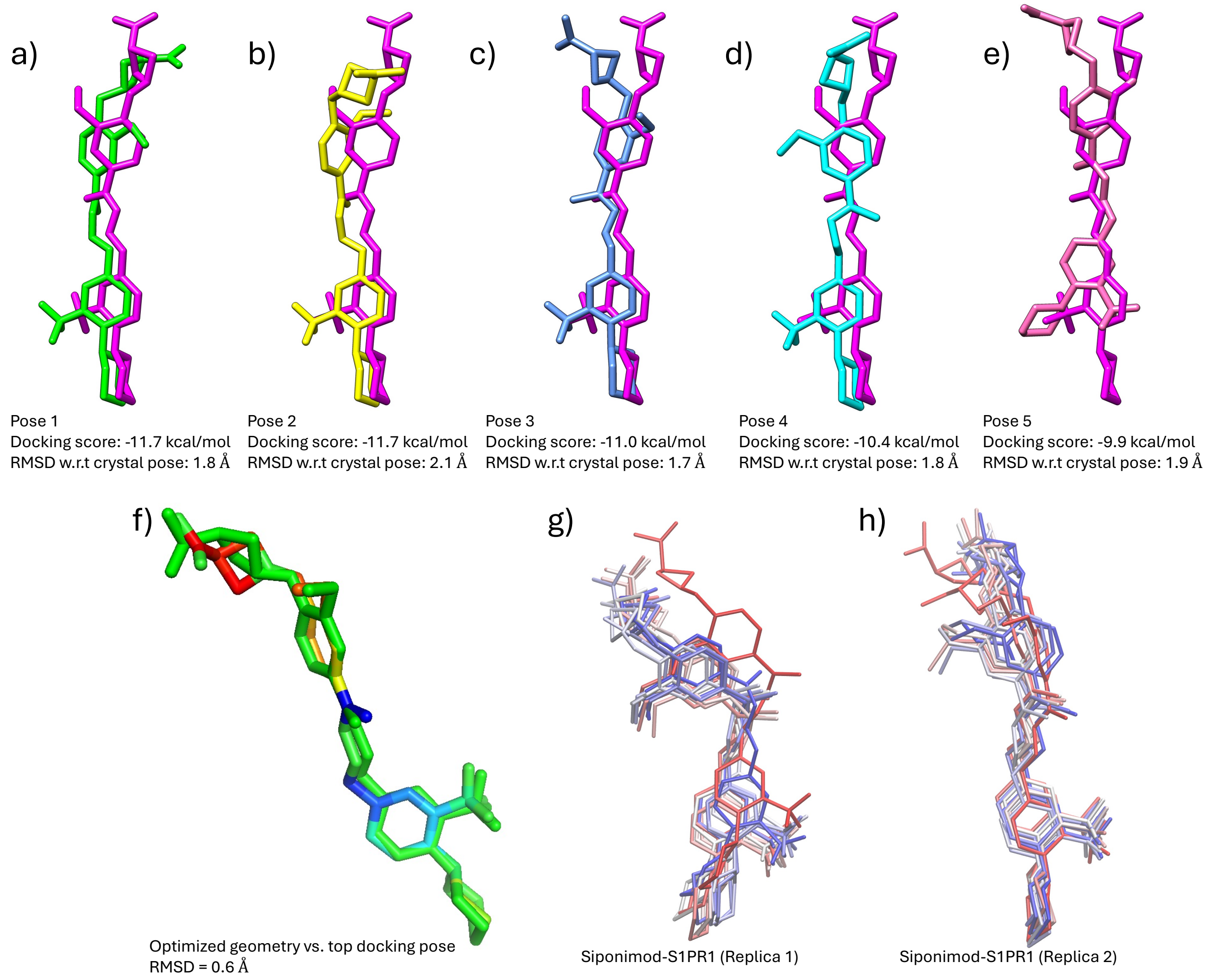

**Figure SI 3.** (a–e) Top five docked poses of Siponimod overlaid with its experimental pose (magenta) from the crystal structure (PDB ID: 7TD4). Symmetry-corrected RMSD values between each docked pose and the experimental pose are indicated. The pose shown in panel (a), having the lowest RMSD, was selected for downstream analyses. (f) Overlay of the top docked pose (green) with the geometry-optimized structure of Siponimod (CPK color scheme) obtained via semi-empirical DFT (xTB) calculations. (g) Representative orientations of Siponimod sampled during replica 1 of the Siponimod–S1PR1 MD simulation (frames shown every 100 ns), overlaid on the experimentally determined structure (red). (h) Representative orientations of Siponimod from replica 2 of the Siponimod–S1PR1 simulation, similarly overlaid on the crystal pose (red).

### 5. Assessing System Fluctuations via RMSD and RMSF Analysis of MD Trajectories

#### Root Mean square Fluctuation

Root Mean Square Fluctuations (RMSF) measures the fluctuations of individual residues averaged over the entire trajectory (T = 1 μs) and depicts their freedom of movement. It is computed using equation 1.

$$RMSF = \sqrt{\frac{1}{T}\sum_{t=1}^{T}(|r_i(t) - r_i^{ref}|)^2} \quad (1)$$

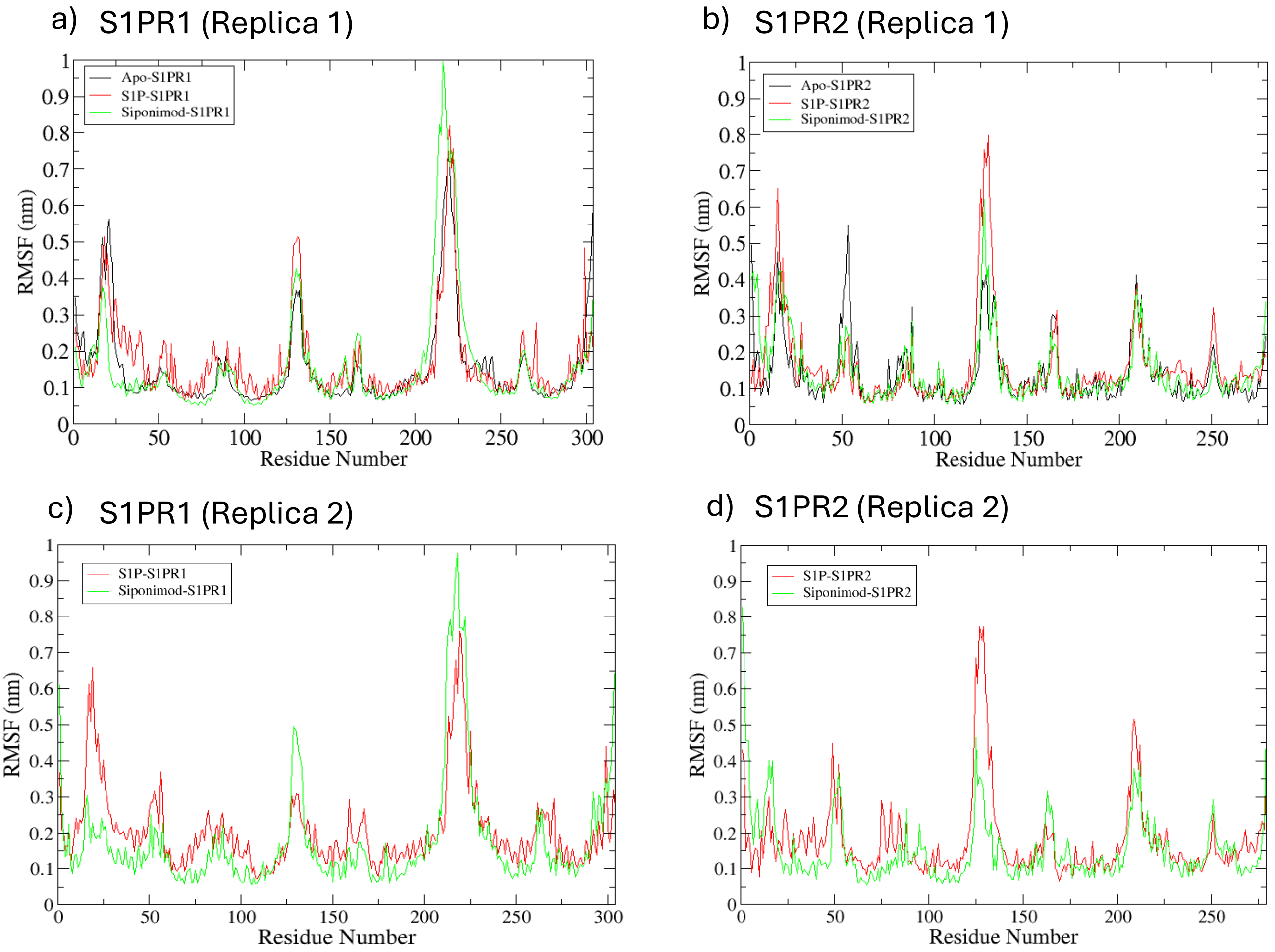

**Figure SI 4.** Residue-wise RMSF values a) of S1PR1 in Apo (black), S1P bound (red) and Siponimod bound (green) state; b) of S1PR2 in Apo (black), S1P bound (red) and Siponimod bound (green) state, as observed in the first replica of the ≈ 1μs long MD trajectory. Residue-wise RMSF values c) of S1PR1 in S1P bound (red) and Siponimod bound (green) state; and d) of S1PR2 in S1P bound (red) and Siponimod bound (green) state, as observed in the second replica of the ≈ 1μs long MD trajectory. High fluctuations corresponds to the residues in intracellular loop regions.

### Root Mean Square Displacement

Root Mean Square Displacement (RMSD) which provides an estimate of the average displacement of each atom ($r_i$) from their initial position ($r_i^{ref}$) over the course of the trajectory (time=t) is calculated using equation 2. Here, $N$ is the total number of atoms on which RMSD is calculated.

$$RMSD = \sqrt{\frac{1}{N}\sum_{i=1}^{N}(r_i(t) - r_i^{ref})^2} \qquad (2)$$

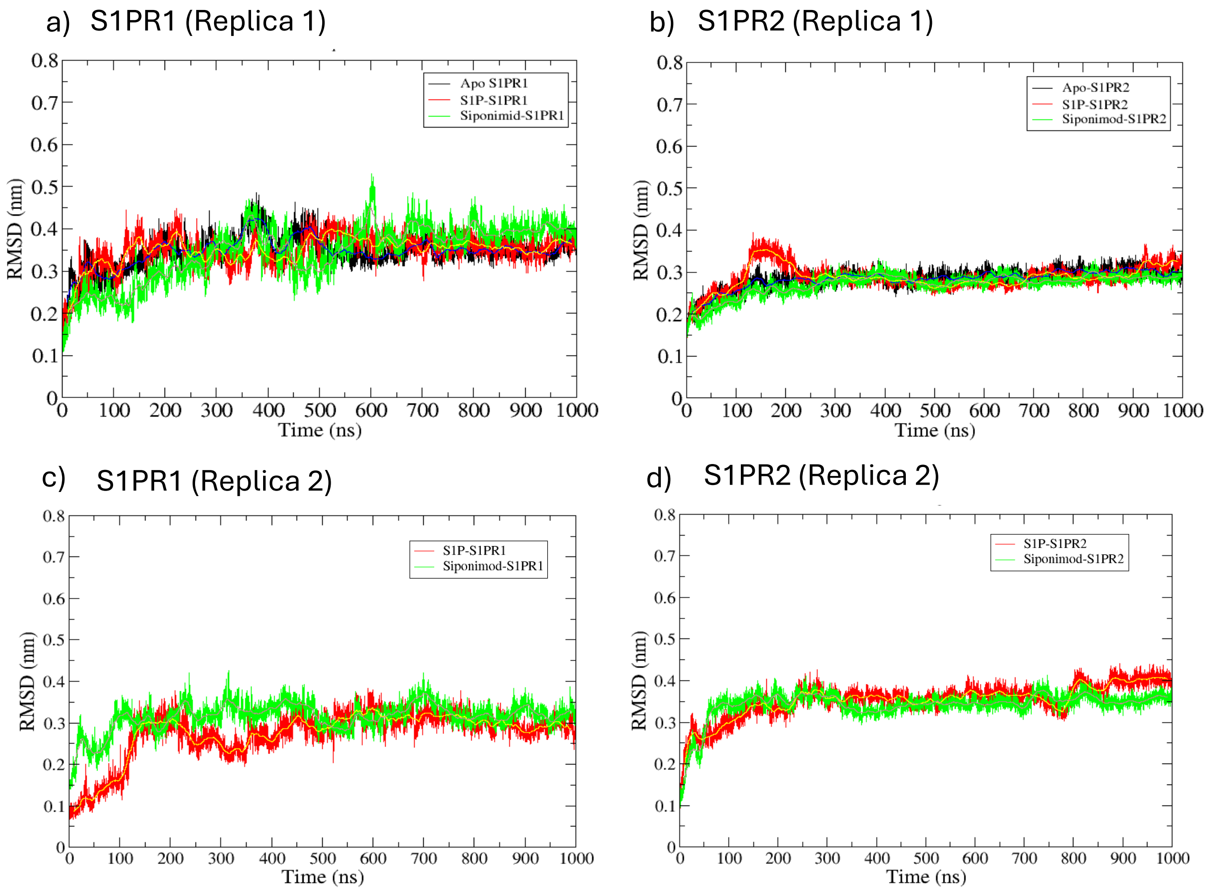

**Figure SI 5.** RMSD fluctuations a) of S1PR1 in Apo (black), S1P bound (red) and Siponimod bound (green) state; b) of S1PR2 in Apo (black), S1P bound (red) and Siponimod bound (green) state as observed in the first replica of the ≈ 1μs long MD trajectory. RMSD fluctuations c) of S1PR1 in S1P bound (red) and Siponimod bound (green) state; d) of S1PR2 in S1P bound (red) and Siponimod bound (green) state, as observed in the second replica of the ≈ 1μs long MD trajectory.

### 6. Principal Component Analysis of the MD trajectories

Principal Component Analysis (PCA)[8] was done using CPPTRAJ in AmberTools22. PCA helps in visualizing the most important macromolecular motions in a reduced dimension. It first creates a covariance matrix for the atomic coordinates of heavy atoms ranked as per the associated weights or the variances to ultimately produce anisotropic atomic movements[9], [10]. Since the first two principal component vectors, PC1 and PC2, are known to capture most of the information about the system's dynamics, we plotted a PC1 vs PC2 scatter plot to visualize how the two independent replicas sample the conformational space.

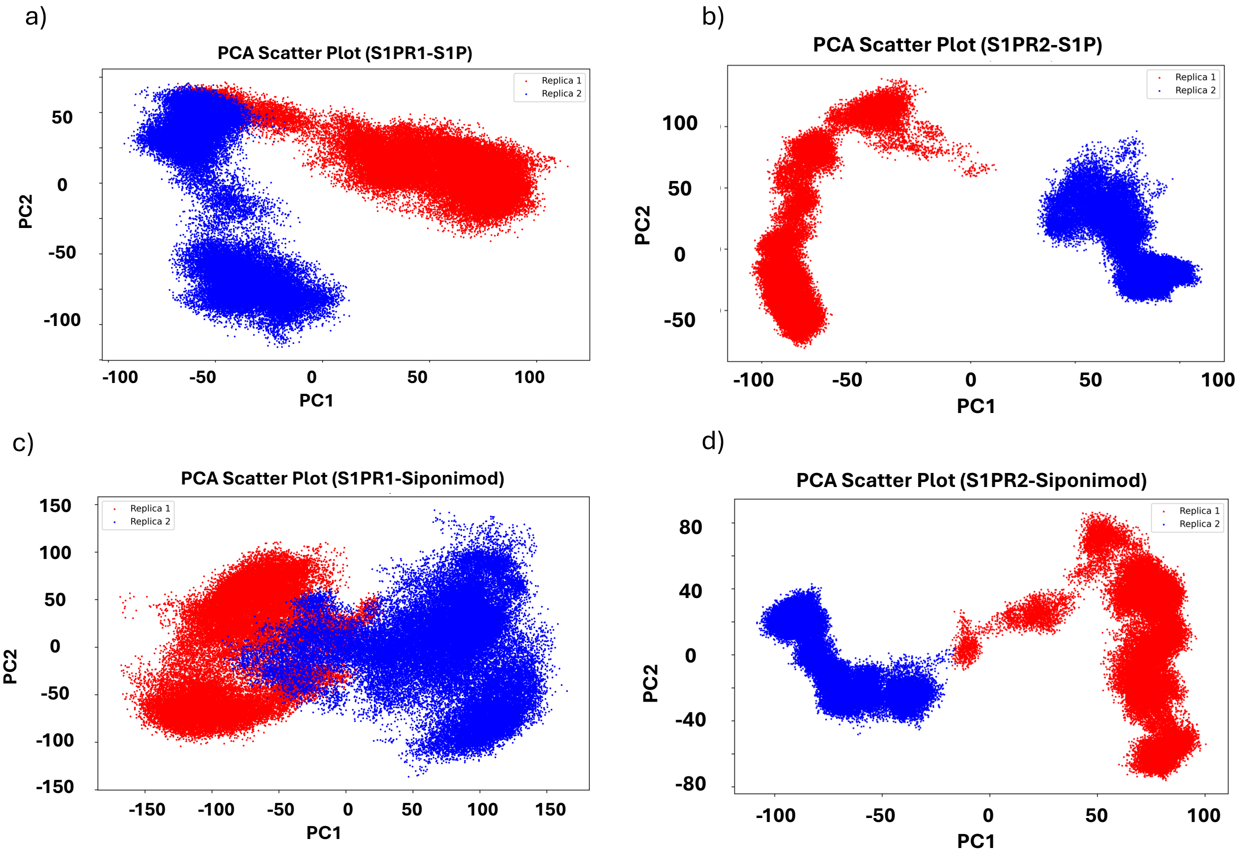

**Figure SI 6.** The conformational space covered by the two independent replicas in the reduces space defined by PC1 and PC2 for a) S1P bound S1PR1, b) S1P bound S1PR2, c) Siponimod bound S1PR1 and d) Siponimod bound S1PR2. Each frame of the first replica are shown in red dots and each frame of the second replica are shown in blue dots.

These anisotropic atomic movements can be visualized through motion vectors where the direction of vectors reflects the direction of motion of the moiety or domain, and the length determines the amplitude of motion. Larger the length of the motion vector, more pronounced is the motion. For that, the reduced motion along the first principal component PC1 (that technically captures the primary collective motions or conformational changes) was studied. As expected, maximum movements take place around the flexible loops regions in the extracellular and intracellular regions in all the systems. To remove additional noise from porcupine graphs for PC1, motion vectors focusing upon TM6 and TM7 are shown (Figure SI 7).

In S1PR1, for both S1P bound and Siponimod bound systems (a) TM7 shows an inwards and (b) bottom part of TM6 around ICL3 shows an outwards motion (Figure SI 7a & Figure SI 7b). In the S1P-S1PR2 system (Figure SI 7c), (a) TM7 moves outward and (b) TM6 exhibits inward movement in the middle section. In contrast, the Siponimod-S1PR2 system (Figure SI 7d) (a) TM6 & TM7 motion vectors moves towards each other.

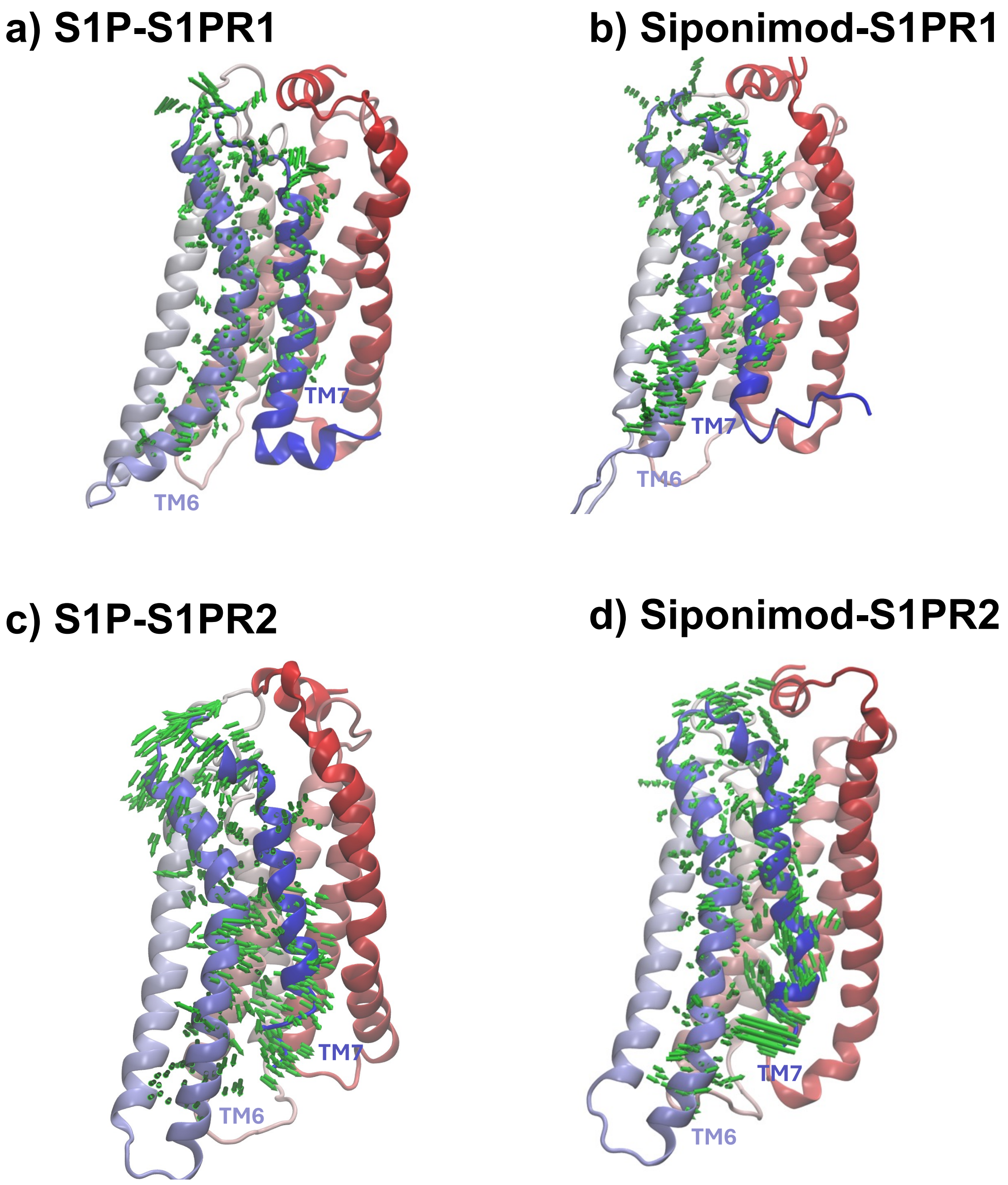

**Figure SI 7.** Porcupine plots illustrating the direction and amplitude of motion along the first principal component (PC1) derived from PCA analysis. Emphasis is placed on the movements of transmembrane helices 6 (TM6) and 7 (TM7), which are known to be critical for GPCR activation. In panels (a, b, and c), TM6 and TM7 tend to move apart near the cytoplasmic region. However, in the case of Siponimod-bound S1PR2 (panel d), the two helices appear to move towards each other.

### 7. Markov State Model using PCA

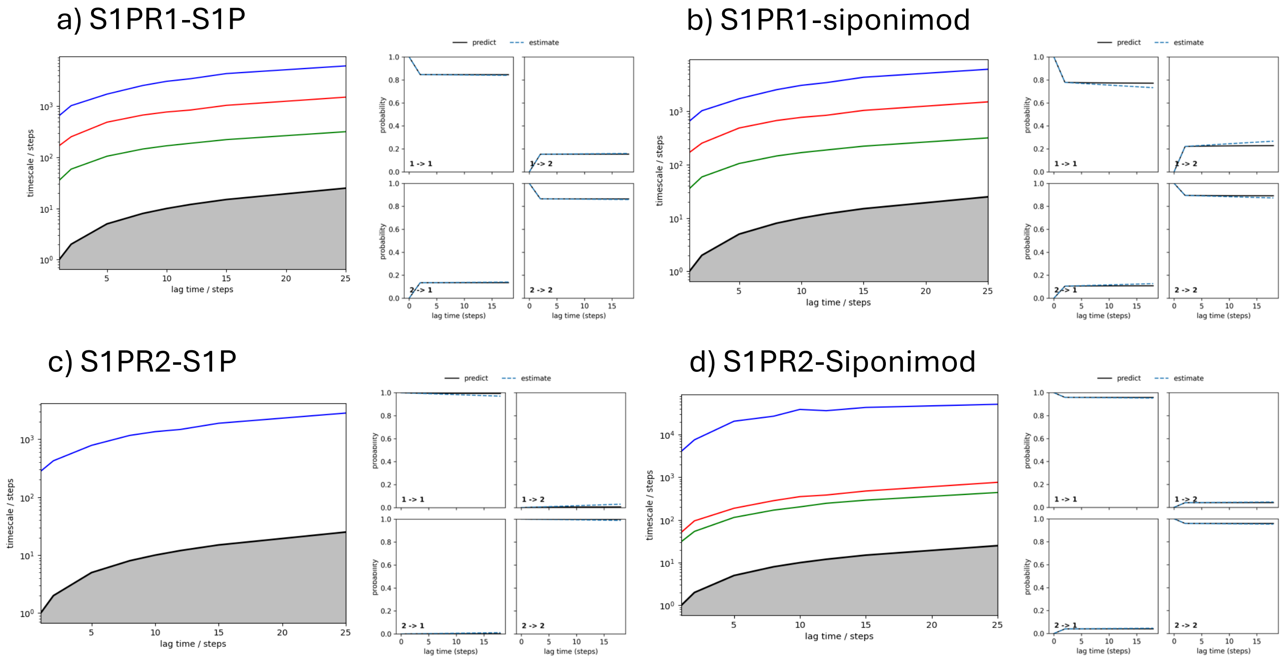

**Figure SI 8.** Implied timescales and Chapman-Kolmogorov (CK) tests for the four simulated systems. The left panels (a–d) show the implied timescale (ITS) plots for each system, based on varying lag times, while the corresponding right panels depict the CK test results using a selected lag time of 20 ns. The good agreement between the estimated (dots) and predicted (lines) transition probabilities in the CK plots confirms the Markovianity of the constructed models at this lag time. Panels (a) and (b) indicate that the kinetic profiles of S1PR1 bound to S1P and Siponimod are highly similar, with comparable timescale hierarchies and excellent CK validation. In contrast, panels (c) and (d) suggest distinct kinetic features in the S1PR2 systems, especially in the Siponimod-bound case, where intermediate timescales diverge slightly, although the model still retains reliable Markovian behaviour overall.

### 8. $\tau$-Random Acceleration Molecular Dynamics (τ-RAMD)

**Table SI 1.** Comparison of residence time obtained from τ-RAMD calculations averaged over 5 replicas of independent MD simulations

| System | Residence time (τ) in ns |
|---|---|
| Siponimod bound S1PR1 | 8.5 |
| Siponimod bound S1PR2 | 18.0 |

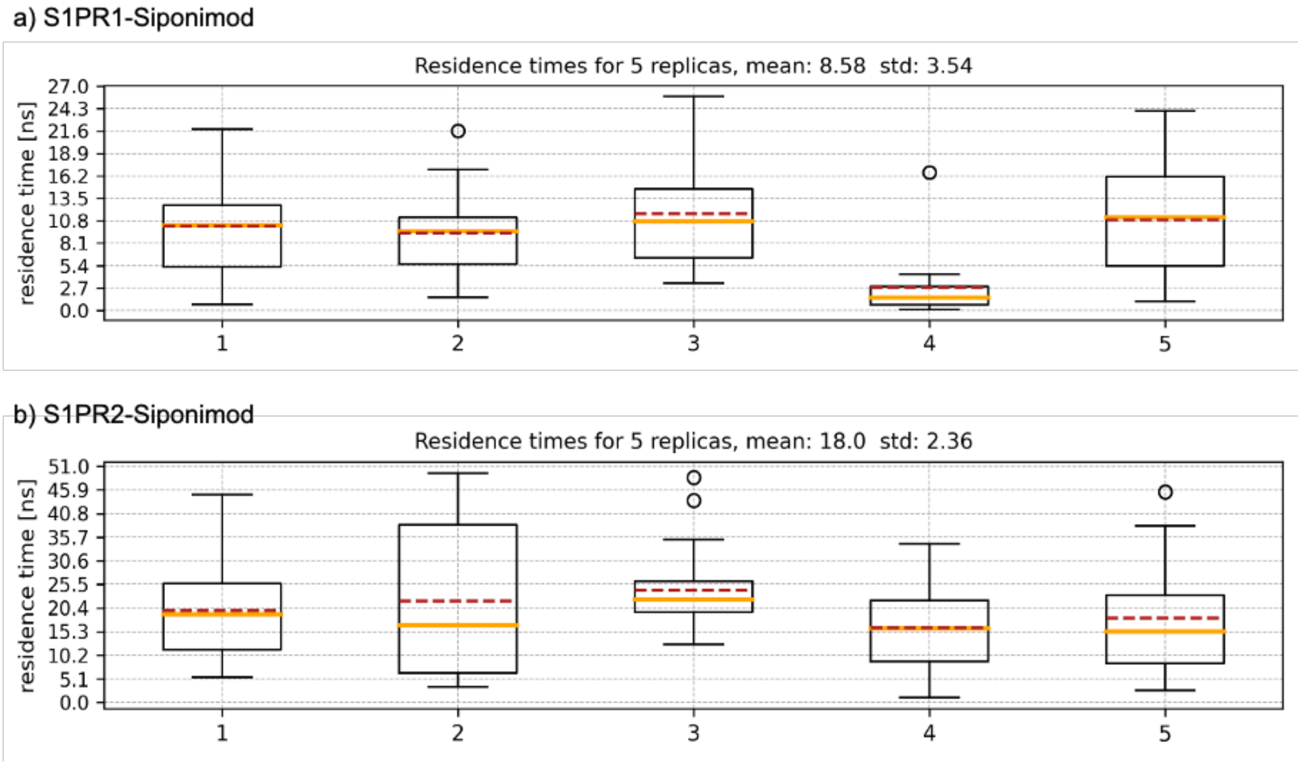

**Figure SI 9.** Estimation of ligand residence times using the τ-RAMD method for Siponimod bound to S1PR1 and S1PR2. Box plots represent residence times (in nanoseconds) computed over five independent τ-RAMD replicas for (a) S1PR1–Siponimod and (b) S1PR2–Siponimod systems. The horizontal dashed red lines indicate the median, and the solid orange lines represent the mean residence time across replicas. Error bars and data spread reflect inter-replica variability. The analysis demonstrates longer average residence times for Siponimod in S1PR2 compared to S1PR1, suggesting stronger kinetic stability in the S1PR2 binding pocket. These results support the reproducibility and robustness of our statistical averaging protocol for residence time estimation.

### 9. Contribution of the Receptor Residues Towards Binding Free Energy of the Ligands (S1P and Siponimod)

Steps followed to consolidate the pair-wise energy decomposition data and identify key residues that contribute consistently to stabilizing S1P and Siponimod at the active sites of S1PR1 and S1PR2.

1. **All interactions that stabilize ligand binding:**

   Shortlist all the interactions that involve the ligand (S1P or Siponimod) from both the replicas

2. **Interactions that are strong:**

   From the cases shortlisted in the previous step, keep only those interactions whose total contribution is 1 kcal/mol or more.

3. **Interactions that are consistent:**

   From the cases shortlisted in the previous step, keep only those interactions that are common in both the replicas

4. **Measuring the mean contributions:**

   Take an average of the energy component values obtained from the two replicas. These values were used in plotting the Figure 3 in main manuscript.

### 10. Analyzing Inter-Residue Dynamics Using Dynamic Cross-Correlation Maps (DCCM)

DCCM between a pair of residues in a protein or protein complex measures the degree to which they are moving together. The DCCM is a NxN heatmap where N represents the number of residues (or C alpha atoms) in the system of interest. The Dynamic Cross Correlation (DCC) between two atoms, $i$ & $j$, can be calculated using equation 3, where $\boldsymbol{r_i}(t)$ represents the vector for atomic coordinates of atom $i$ over ensemble average.

$$DCC(i,j) = \frac{\langle \Delta \boldsymbol{r_i}(t) . \Delta \boldsymbol{r_j}(t) \rangle}{\left( \sqrt{\langle \Delta \mathbf{r}_i^2(t) \rangle} \sqrt{\langle \Delta \mathbf{r}_j^2(t) \rangle} \right)} \quad (3)$$

S1PR1

| RESIDUE ID (shifted) | Transmembrane Helix |
|---|---|
| 26-47 | TM1 |
| 62-83 | TM2 |
| 96-117 | TM3 |
| 140-161 | TM4 |
| 176-203 | TM5 |
| 237-257 | TM6 |
| 269-289 | TM7 |

| RESIDUE ID (shifted) | Outside Membrane |
|---|---|
| 1-25 | NT |
| 48-61 | ICL1 |
| 84-95 | ECL1 |
| 118-139 | ICL2 |
| 162-175 | ECL2 |
| 204-236 | ICL3 |
| 258-268 | ECL3 |
| 290-361 | CT |

S1PR2

| RESIDUE ID (shifted) | Transmembrane Helix |
|---|---|
| 24-48 | TM1 |
| 56-84 | TM2 |
| 99-117 | TM3 |
| 137-162 | TM4 |
| 179-199 | TM5 |
| 223-244 | TM6 |
| 261-281 | TM7 |

| RESIDUE ID (shifted) | Outside Membrane |
|---|---|
| 1-23 | NT |
| 49-55 | ICL1 |
| 85-98 | ECL1 |
| 118-136 | ICL2 |
| 163-178 | ECL2 |
| 200-222 | ICL3 |
| 245-260 | ECL3 |
| 282-342 | CT |

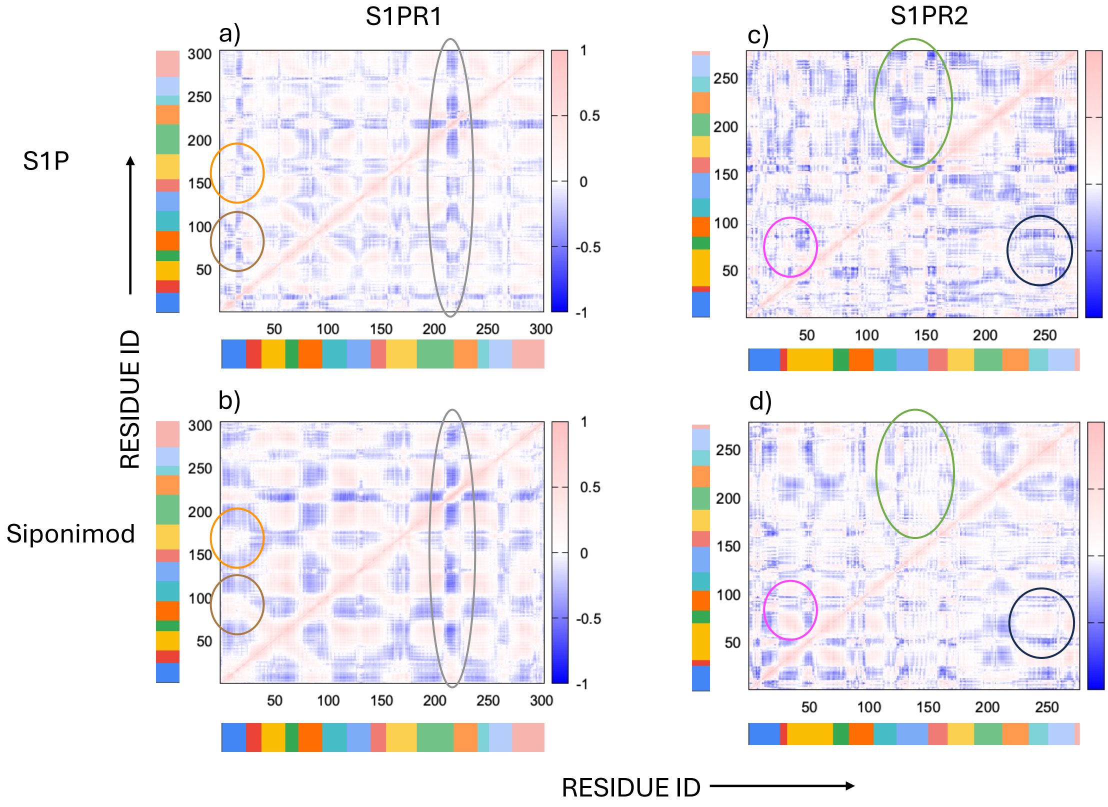

**Figure SI 10.** DCCM for S1PR1 protein in (a) S1P bound and (b) Siponimod bound states. Same for S1PR2 protein in (c) S1P bound and (d) Siponimod bound states. Residue mapping to NT, ECL1-3, ICL1-3, CT and TM1-7 shown in the tables above. The colour gradient starts with blue, representing anti-correlated motion (-1), transitions through white to indicate no correlation (0), and gradually shifts to pink, illustrating highly correlated motion (+1) between residues. Key differences between the S1P-bound and Siponimod-bound receptors are highlighted in circles using distinct colours : brown – ECL1 vs. NT in S1PR1, orange – ECL2 vs. NT in S1PR1, grey – ICL3 vs. rest of S1PR1, magenta – TM2 vs. ICL1 in S1PR2, black - TM2 vs TM6 in S1PR2, green – ICL2 vs (TM4-ECL2-TM5-ICL3-TM6-ECL3-TM7) in S1PR2

### 11. Changes in Transmission Switches on Ligand Binding in S1PR1 and S1PR2

#### Transmission switches in S1PR1

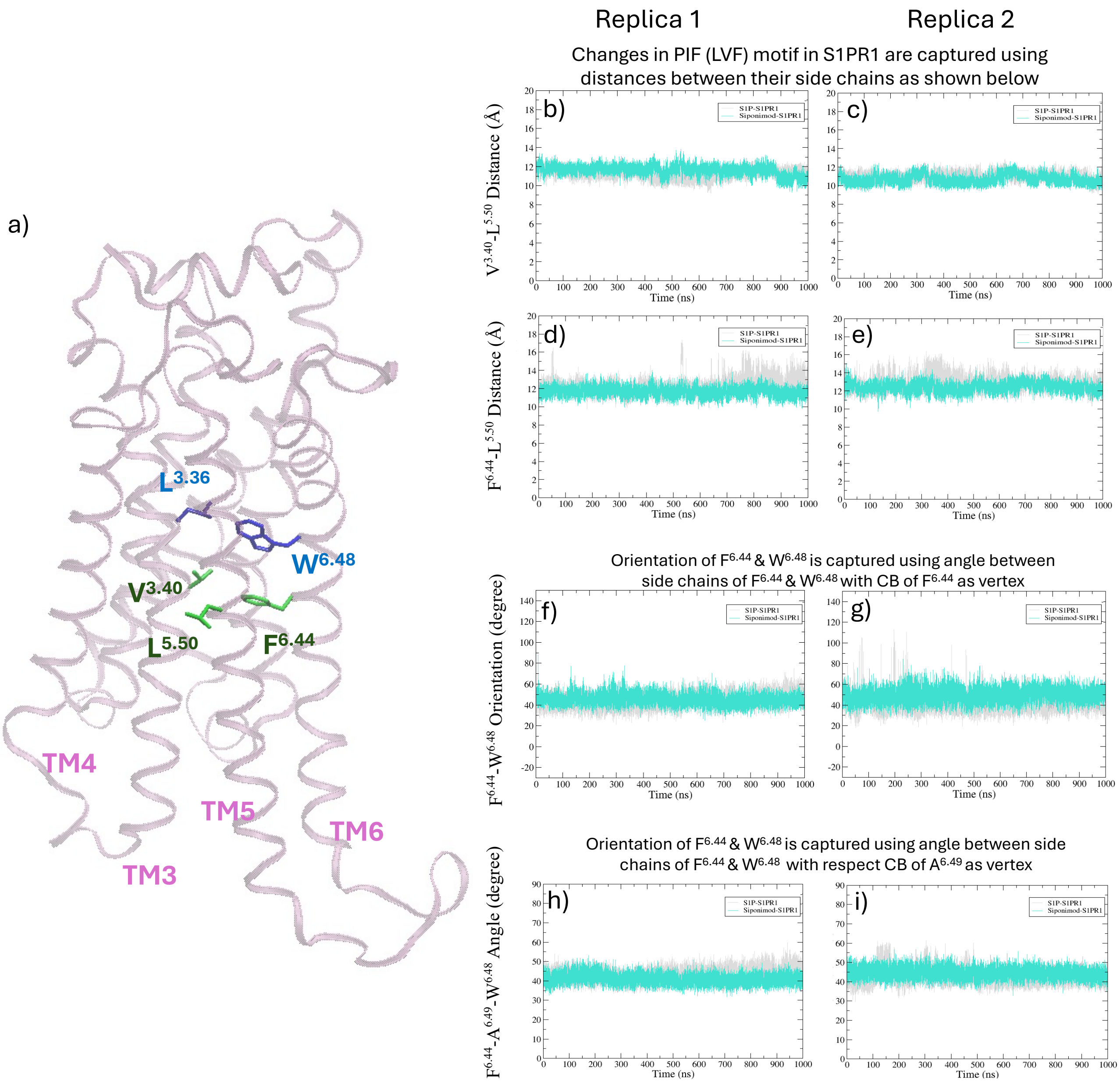

**The caption of the above image is provided in the next page**

**Figure SI 11.** Tracking the variation in transmission switches in S1PR1 during the simulation through customized collective variables (CVs). (a) Cartoon representation of the backbone of S1PR1. The PIF (LVF) motif is shown in green sticks. The FW switch is shown in purple sticks. Values for the S1P-bound and Siponimod-bound systems are plotted using grey and turquoise colour, respectively in panel b-i . Two different CVs were used to track the changes in the PIF(LVF) motif: the distance between $L^{5.50}$ & $V^{3.40}$ (panel b and c, for the first and the second replica respectively), and angle between side chains of $F^{6.44}$ & $W^{6.48}$ with atom CB of $F^{6.44}$ as vertex (panel d and e, for the first and the second replica respectively). Two different CVs were used to track the changes in the orientation of the FW switch: angle between side chains of $F^{6.44}$ & $W^{6.48}$ with atom CB of $F^{6.44}$ as vertex (panel f and g, for the first and the second replica respectively), and angle between side chains of $F^{6.44}$ & $W^{6.48}$ with atom CB of $A^{6.49}$ as vertex(panel h and i, for the first and the second replica respectively). Both S1P and Siponimod bound S1PR1 systems are showing a constant fluctuation indicating the starting active state of the receptor is maintained by the transmission switches.

### Transmission switches in S1PR2

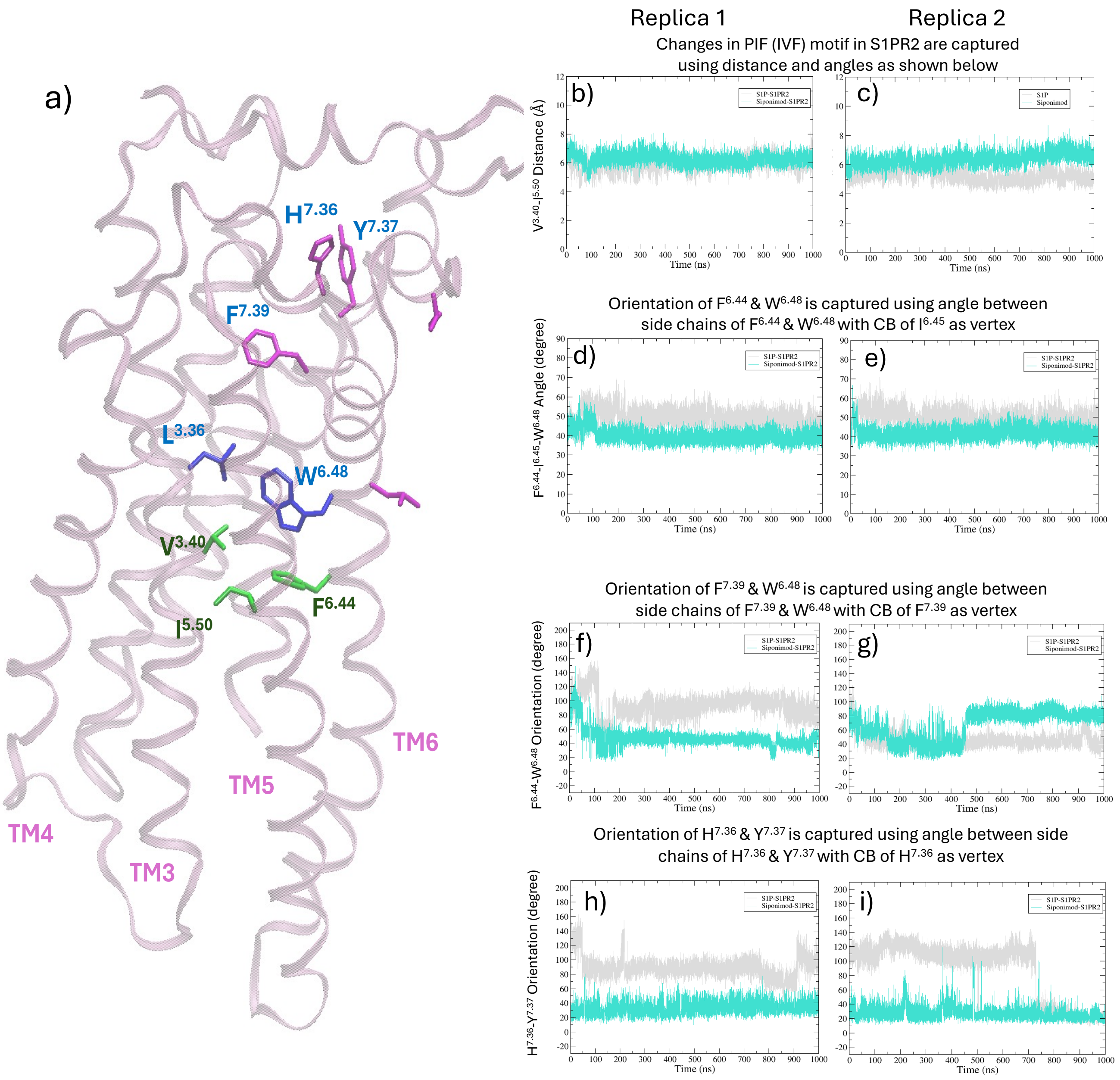

**The caption of the above image is provided in the next page**

**Figure SI 12.** Tracking the variation in transmission switches in S1PR2 during the simulation through customized collective variables (CVs). (a) Cartoon representation of the backbone of S1PR1. The PIF (IVF) motif is shown in green sticks. The FW switch is shown in purple sticks. Other residues showing crucial conformational changes are shown in magenta sticks. Values for the S1P-bound and Siponimod-bound systems are plotted using grey and turquoise colour, respectively in panel b-i. The distance between $I^{5.50}$& $V^{3.40}$ is used as a CV to track changes in the in the PIF (IVF) motif. The distance is maintained during the first replica (panel b) but a difference of ~2 Å between S1P and Siponimod bound receptors is seen towards end of trajectory in the second replica (panel c). The orientation of $F^{6.44}$ & $W^{6.48}$ is captured using angle between side chains of $F^{6.44}$ & $W^{6.48}$ with CB of $I^{6.45}$ as vertex (panel d and e, for the first and the second replica respectively). A slight difference of 10° between S1P and Siponimod bound cases is evident post 100 ns. In S1P bound active conformation $F^{6.44}$ tends to stay outwards. The orientation of $F^{7.39}$ & $W^{6.48}$ is captured using angle between side chains of $F^{7.39}$ & $W^{6.48}$ with atom CB of $F^{7.39}$ as vertex (panel f and g, for the first and the second replica respectively). The trend for both S1P and Siponimod bound S1PR2 seems different in 2 replicas. The orientation of $H^{7.36}$ & $Y^{7.37}$ is captured using angle between side chains of $H^{7.36}$ & $Y^{7.37}$ with CB of $H^{7.36}$ as vertex (panel h and i, for the first and the second replica respectively). While the orientation between two residues is frozen in Siponimod bound S1PR2 conformation, it is changing frequently in case of S1P bound S1PR2.

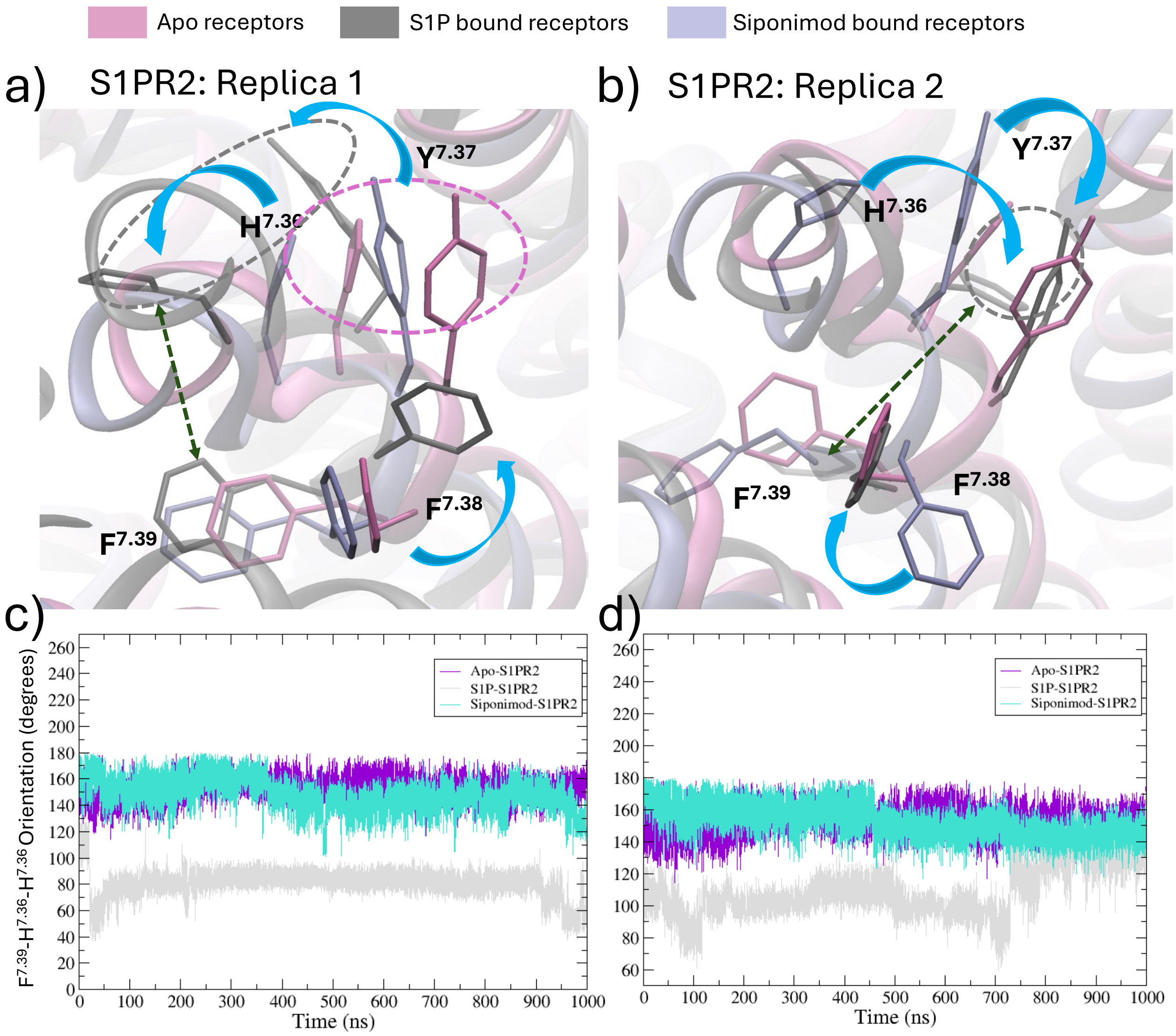

**Figure SI 13.** Side-chain conformations of residues in the extracellular region of TM7 in S1PR2.
(a) Conformational arrangement of residues $H^{7.36}$, $Y^{7.37}$, $F^{7.38}$, and $F^{7.39}$ in Replica 1 for the Apo (mauve), S1P-bound (grey), and Siponimod-bound (ice-blue) S1PR2 systems.
(b) Same set of residues shown for Replica 2 of the S1P- and Siponimod-bound systems. For the Apo system, data from Replica 1 is reused, as only one replica was generated.
(c–d) Relative orientations of key residues $H^{7.36}$ and $F^{7.39}$ are highlighted for Replica 1 (panel c) and Replica 2 (panel d). Despite the differences in sampling between the two independent replicas, the relative orientations of $H^{7.36}$ and $F^{7.39}$ remain consistent across S1P- and Siponimod-bound S1PR2 systems.

### 12. Validation of conformational switching through additional replicates of S1P & Siponimod bound S1PR2

We carried out one additional 500 ns replicate each for the S1P- and Siponimod-bound S1PR2 systems to observe if the crucial switches orient differently in S1P vs. Siponimod bound systems, as observed in the first two replicates. The newly generated trajectories reaffirm our earlier findings: upon Siponimod binding, the two key transmission switches—specifically the aromatic residues in the extracellular segment of TM7—adopt a distinct conformational configuration compared to the S1P-bound state. Furthermore, the relative displacement and angular orientation of TM7 with respect to TM6 also mirror the trends seen in the initial replicas. Together, these results strengthen our conclusion that Siponimod binding induces a unique conformational response in S1PR2, one that differs markedly from the canonical activation pattern triggered by S1P.

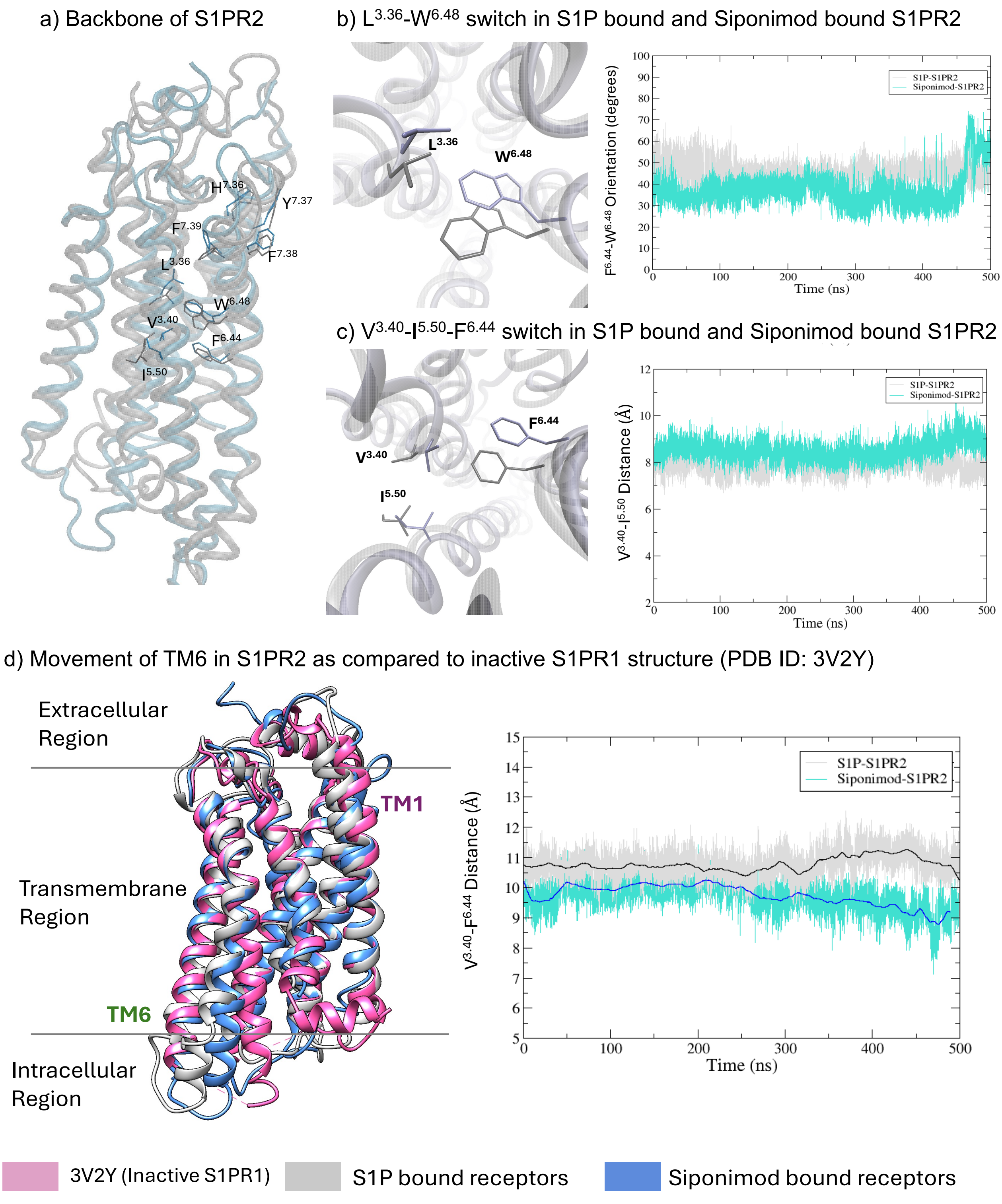

**Figure SI 14.** (a) Backbone of S1P bound S1PR2 superposed over Siponimod bound S1PR2 with key residues constructing the $L^{3.36}$-$W^{6.48}$ and $V^{3.40}$-$I^{5.50}$-$F^{6.44}$ switches labeled. (b) Comparison of the S1P bound and Siponimod bound conformation of the $L^{3.36}$-$W^{6.48}$ switch, with the changes in appropriate CV with simulation time. (c) Comparison of the S1P bound and Siponimod bound conformation of the $V^{3.40}$-$I^{5.50}$-$F^{6.44}$ switch, with the changes in appropriate CV with simulation time. (d) Movement of the TM6 helix towards inactive state (shown in pink) on Siponimod binding (shown in ice blue) from the S1P bound active state (shown in gray). The same movement tracked using change in the appropriate CV with simulation time.

### 13. Details of Collective Variables (CVs) used for analysis of MD trajectories

**Table SI 2:** Details of CVs used for analysing the MD simulation trajectories of the ligand free and ligand-bound systems.

| Figure | CV name | Unit | Atoms used for measurement |
|---|---|---|---|
| Figure 5 | $V^{3.40}$-$F^{6.44}$ Distance | Å | CB($V^{3.40}$)-CB($F^{6.44}$) |
| Figure SI 10b, 10c | $V^{3.40}$-$L^{5.50}$ Distance | Å | CB($V^{3.40}$)-CB($L^{5.50}$) |
| Figure SI 10d, 10e | $F^{6.44}$-$L^{5.50}$ Distance | Å | CZ($F^{6.44}$)-CG($L^{5.50}$) |
| Figure SI 10f, 10g | $F^{6.44}$-$W^{6.48}$ Orientation | degree | CZ($F^{6.44}$)-CB($F^{6.44}$)-CZ3($W^{6.48}$) |
| Figure SI 10h, 10i | $F^{6.44}$-$A^{6.49}$-$W^{6.48}$ Angle | degree | CB($F^{6.44}$)-CB($A^{6.49}$)-CB($W^{6.48}$) |
| Figure SI 11b, 11c | $V^{3.40}$-$I^{5.50}$ Distance | Å | CB($V^{3.40}$)-CB($I^{5.50}$) |
| Figure SI 11d, 11e | $F^{6.44}$-$I^{6.45}$-$W^{6.48}$ Angle | degree | CB($F^{6.44}$)-CB($I^{6.45}$)-CB($W^{6.48}$) |
| Figure SI 11f 11g | $F^{7.39}$-$W^{6.48}$ Orientation | degree | CZ($F^{7.39}$)-CB($F^{7.39}$)-CZ3($W^{6.48}$) |
| Figure SI 11h, 11i | $H^{7.36}$-$Y^{7.37}$ Orientation | degree | CE1($H^{7.36}$)-CB($H^{7.36}$)-CZ($Y^{7.37}$) |
| Figure SI 12 | $F^{7.39}$-$H^{7.36}$ Orientation | degree | NE2($F^{7.39}$)-CB($H^{7.36}$)-CZ($H^{7.36}$) |
